# Major diversification of caribou shaped by glacial cycles before the Last Glacial Maximum

**DOI:** 10.1101/2020.11.30.404731

**Authors:** Rebecca S. Taylor, Micheline Manseau, Cornelya F. C. Klütsch, Jean L. Polfus, Audrey Steedman, Dave Hervieux, Allicia Kelly, Nicholas C. Larter, Mary Gamberg, Helen Schwantje, Paul J. Wilson

## Abstract

Pleistocene glacial cycles influenced the diversification of high-latitude wildlife species through recurrent periods of range contraction, isolation, divergence, and expansion from refugia and subsequent admixture of refugial populations. For many taxa, research has focused on genetic patterns since the Last Glacial Maximum (LGM), however glacial cycles before the LGM likely impacted genomic variation which may influence contemporary genetic patterns. We investigate diversification and the introgressive history of caribou (*Rangifer tarandus*) in western Canada using 33 high-coverage whole genomes coupled with larger-scale mitochondrial data. Contrary to the well-established paradigm that caribou ecotypes and contemporary genetic diversity arose from two major lineages in separate refugia during the LGM, a Beringian-Eurasian (BEL) and a North American (NAL) lineage, we found that the major diversifications of caribou occurred much earlier at around 110 kya, the start of the last glacial period. Additionally, we found effective population sizes of some caribou reaching ~700,000 to 1,000,000 individuals, one of the highest recorded historical effective population sizes for any mammal species thus far. Mitochondrial analyses dated introgression events prior to the LGM dating to 20-30 kya and even more ancient at 60 kya, coinciding with colder periods with extensive ice coverage, further demonstrating the importance of glacial cycles and events prior to the LGM in shaping demographic history. Reconstructing the origins and differential introgressive history has implications for predictions on species responses under climate change. Our results highlight the need to investigate pre-LGM demographic patterns to fully reconstruct the origin of species diversity, especially for high-latitude species.

## 1| INTRODUCTION

The evolution of terrestrial flora and fauna was strongly impacted by glacial fluctuations during the Pleistocene (Hewitt, 2004; Shafer, Cullingham, Côté, & Coltman, 2010; Soltis, 2006) that caused repeated range shifts and restricted populations to ice-free refugia with viable habitat (Hewitt, 2004; Shafer et al., 2010; Soltis, 2006). Glacial cycles have had major biogeographical consequences and have contributed to the diversification of biodiversity by partitioning genetic variation across the landscape through bottlenecks, genetic drift, and mutations that result in genetic differentiation and local adaptations in geographically separated refugia (Hewitt, 2004).

Earlier work suggests that genetic diversity in western North American wildlife populations is the result of recurrent cycles of range shifts caused by expansion and contraction, isolation, and potential introgression that, in combination with habitat heterogeneity, potentially explains the observed intraspecific diversity and local adaption (Campbell, Takebayashi, & López, 2015; Galbreath, Cook, Eddingsaas, & DeChaine, 2011; Shafer et al., 2010). However, studies on many taxa have focused on the Last Glacial Maximum (LGM; 26.5-19 kya), and events since then, in reconstructing the evolutionary history of wildlife species. For example, studies of demographic expansion in downy woodpeckers (*Picoides pubescens*; Pulgarín-R & Burg, 2012), colonization history and genetic variation in muskox (*Ovibos moschatus*; Hansen et al., 2018), dispersal routes and bison phylogeography (*Bison* sp.; Heintzman et al., 2016), and subspecies and genetic diversity of tree species in western North America (Roberts & Hamann, 2015) all provided valuable information on how the LGM and more recent times shaped phylogeography but did not investigate the impact of events prior to the LGM. However, population sizes have been shown to fluctuate considerably throughout the Quaternary, such as in a demographic reconstruction of 38 avian species (Nadachowska-Brzyska, Smeds, Zhang, & Ellegren, 2015), and of 11 bat species (Chattopadhyay, Garg, Ray, & Rheindt, 2019), among others (e.g. Kozma, Melsted, Magnússon, & Höglund, 2016; Louis et al., 2020; Miller et al., 2012). Multiple cycles of population increase and decline, and associated range shifts, are likely to have influenced the processes of adaptation and diversification. Further studies of temporal dynamics during different glacial cycles will help us to understand the impact of climate change on biodiversity (Nadachowska-Brzyska et al., 2015; Kozma et al., 2016).

In north-western North America, caribou (*Rangifer tarandus*) comprise two genetic lineages which are thought to have evolved around 120 kya (Banfield, 1961; Polfus, Manseau, Klütsch, Simmons, & Wilson, 2017; Taylor et al., 2020). However, most genetic studies have focussed on the impacts of the LGM, and events since then, on ecotype evolution, demographic history, and contemporary genetic diversity. The two lineages were in separate refugia during the LGM: the Beringian-Eurasian lineage (BEL) in the Beringian refugium, and the North American lineage (NAL) were south of the ice sheets (Klütsch, Manseau, & Wilson, 2012; McDevitt et al., 2009; Weckworth, McDevitt, Musiani, Hebblewhite, & Mariani, 2012; Yannic et al., 2013). Caribou recolonized the ice-free landscape after the LGM and experienced admixture upon secondary contact (Klütsch, Manseau, Trim, Polfus, & Wilson, 2016), which has been referred to as a ‘hybrid swarm’ (McDevitt et al., 2009). The fact that the lineages arose much earlier, however, suggests that the well-established paradigm of ecotype evolution during the LGM, followed by admixture within the mountains, may be an oversimplification of the true demographic and introgressive history as drivers of contemporary genetic structure of caribou in north-western North America. We provide an improved demographic history reconstruction using 33 whole-genomes and over 1,800 mitochondrial control region sequences to date the diversification of caribou and introgression events, understand the evolution of caribou ecotypes, and reveal the effects of different glacial periods on contemporary caribou diversity.

North-western North American caribou comprise several contemporary subspecies and Designatable Units (Figure 1; Banfield, 1961; COSEWIC, 2011). The three subspecies found in the region are Grant’s caribou (*Rangifer tarandus granti*), barren-ground caribou (*R. t. groenlandicus*), and woodland caribou (*R. t. caribou*). Grant’s caribou and barren-ground caribou are found in Alaska and northern Canada and are proposed to be of Beringian origin (BEL) whereas most woodland caribou ecotypes are proposed to have evolved south of the ice sheets (NAL) and are distributed throughout the boreal forest as well as the mountain regions (Banfield, 1961, COSEWIC, 2011). However, in caribou, subspecific designations are insufficient to characterize the morphological, behavioural, and ecological variation observed across the species range. To alleviate the subspecific issue for Canadian species, the Committee on the Status of Endangered Wildlife in Canada (COSEWIC, 2011) introduced the Designatable Unit (DU) concept whereby DUs must be able to demonstrate attributes making them both discrete and evolutionarily significant based on the best available evidence.

**FIGURE 1.**
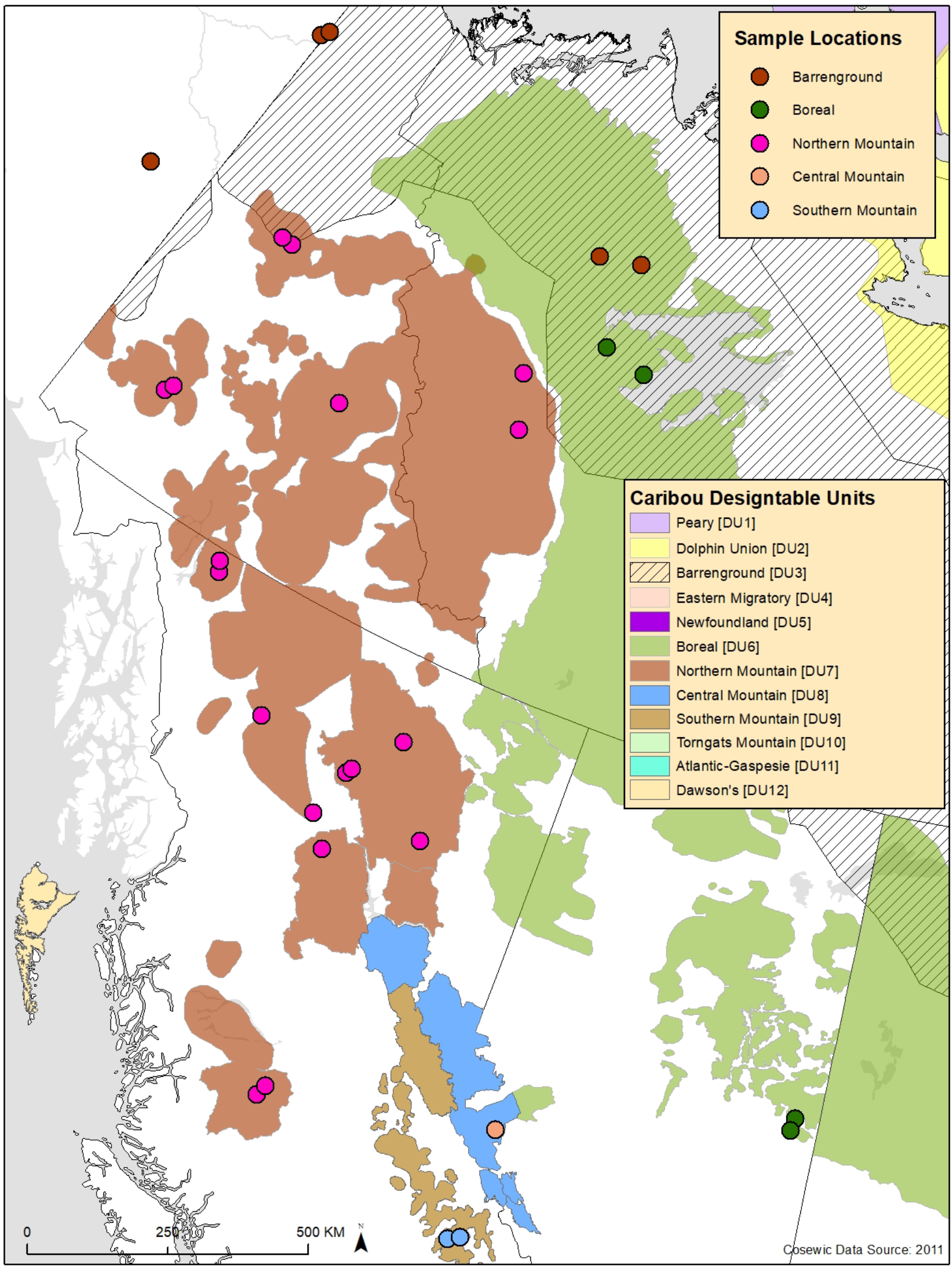
Map of sampling locations of the caribou whole-genome sequences in northwestern North America. Background colours show the ranges of the Canadian Designatable Units and points show the sample locations for the whole genome sequences.

COSEWIC has identified and assessed five DUs in western Canada: barren-ground, boreal, northern mountain, southern mountain, and central mountain (Figure 1; COSEWIC, 2011). Barren-ground caribou (DU3) are typically found in large groups and migrate long distances to calve whereas boreal caribou (DU6) are dispersed throughout the boreal forest in small groups and spread out to calve (COSEWIC, 2011). Boreal caribou are generally considered to belong to the woodland subspecies and are more sedentary than barren-ground caribou. However, caribou that show typical boreal caribou sedentary behaviour in central Northwest Territories belong to the BEL, suggesting parallel phenotypic evolution (Horn et al., 2018; Polfus et al., 2017; Taylor et al., 2020). Northern mountain caribou (DU7) are found in the Yukon, western Northwest Territories, and in northern and central British Columbia. They migrate shorter distances than barren-ground caribou, generally using higher elevation plateaus for calving in the spring/summer and, in some cases, lower elevation valleys in the winter. Northern mountain caribou are generally found in smaller groups than barren-ground caribou (COSEWIC, 2011). Previous studies indicated that northern mountain caribou belong to the BEL and are also considered to be in the woodland caribou subspecies (Polfus et al., 2017; Taylor et al., 2020; Weckworth et al., 2012). Central mountain (DU8) and southern mountain (DU9) caribou are found in the rugged mountains of west-central Alberta and southeast/south-central British Columbia (Klütsch et al., 2012; Weckworth et al., 2012; Weckworth, Hebblewhite, Mariani, & Musiani, 2018). Central mountain caribou also undertake elevational migrations to the eastern slopes of the Rocky Mountains to calve (COSEWIC, 2011). Southern mountain caribou undertake as many as four altitudinal migrations a year, and are associated with old growth conifer forests (COSEWIC, 2011).

We use mitochondrial and whole-genome sequences representing all three subspecies of caribou in north-western North America and the five DUs of western Canada to assess impact of glacial events and climate changes pre-LGM, as well as the LGM, on diversification of the species, the genomic structure, and patterns of introgression. We aim to: (1) accurately date when the diversification of caribou occurred and whether we see introgression events pre-LGM, and (2) investigate whether genomic structure reveals substructuring consistent with a history more complex than two major refugia and subsequent colonization after the LGM. We compare our results to historical patterns of temperature and ice coverage to understand how climate cycles impacted the distribution and abundance of caribou populations.

## 2| MATERIALS AND METHODS

### 2.1| Sample collection and DNA extraction

Caribou tissue and faecal samples were collected by the Government of British Columbia, the Government of Alberta, the Government of the Northwest Territories, communities of the Sahtú region of the Northwest Territories, Parks Canada, Environment and Climate Change Canada, and the Northern Contaminants Program between 1999 and 2018 (complemented by GenBank sequences; Klütsch et al., 2012; Klütsch et al., 2017; McDevitt et al., 2009; Polfus et al., 2017; Roffler, Adams, Talbot, Sage, & Dale, 2012; Weckworth et al., 2012).

For whole genome analysis, we used 18 genomes sequenced for a previous study (Taylor et al., 2020). We combined these with 15 new whole genome sequences, 12 of which were from tissue samples and 3 of which were from faecal samples (Figure 1; Table 1). Two of the tissue and the three faecal genomes were used for a methods paper describing the success of reconstructing genomes using non-invasive samples (Taylor, Manseau, Redquest, & Wilson, 2020b). Tissue samples were extracted using a Qiagen DNAeasy tissue extraction kit following the manufacturer’s instructions (Qiagen, Hilden, Germany). Samples were run on a Qubit fluorometer (Thermo Fisher Scientific, MA, USA) using the High Sensitivity Assay Kit and normalized to 20ng/μl at a final volume of 50μl. The laboratory protocol for faecal samples is described in detail in Taylor et al. (2020b). The DNA was shipped to The Centre for Applied Genomics (TCAG) at the Hospital for Sick Children (Toronto, Ontario) for library preparation and sequencing. The 15 samples, alongside one additional sample not included here, were run on eight lanes of an Illumina HiSeq X (Illumina, San Diego, CA, USA). Raw reads from Taylor et al. (2020) are available on the National Centre for Biotechnology (NCBI) under BioProject accession number PRJNA634908. All new raw reads will be made available at the NCBI upon acceptance.

**TABLE 1.** Information for each caribou genome including sampling location and Individual ID, Designatable Unit (DU), subspecies, collection year and tissue type, whether the sample is new for this manuscript, mean depth of coverage from the BAM file, mean depth in the VCF file, and individual inbreeding co-efficients.

| Location and Individual | DU | Subspecies | Collection Year | Type | New for this study | Mean depth (VCF file) | Inbreeding co-efficient, F |
| --- | --- | --- | --- | --- | --- | --- | --- |
| Northwest Territories, Redstone 15460 | Northern mountain | Woodland | 2013 | Muscle | No | 40.129 | -0.064 |
| Northwest Territories Sahtú region 17825 | Boreal | Woodland | 2013 | Muscle | No | 38.827 | -0.016 |
| Northwest Territories, Redstone 17896 | Northern mountain | Woodland | 2014 | Muscle | No | 37.190 | -0.055 |
| Manitoba Qamanirjuaq 21332 | Barren-ground | Barren-ground | 2008 | Hide | No | 36.978 | -0.073 |
| Manitoba Qamanirjuaq 21350 | Barren-ground | Barren-ground | 2008 | Hide | No | 36.452 | -0.075 |
| Alberta Cold Lake 24461 | Boreal | Woodland | 2014 | Faecal | No | 8.382 | -0.260 |
| Alberta Cold Lake 24476 | Boreal | Woodland | 2014 | Faecal | No | 14.403 | 0.012 |
| Northwest Territories Bluenose 27177 | Barren-ground | Barren-ground | 2013 | Muscle | No | 39.801 | -0.065 |
| Northwest Territories Bluenose 27186 | Barren-ground | Barren-ground | 2013 | Muscle | No | 37.766 | -0.069 |
| Yukon Aishihik 27601 | Northern mountain | Woodland | 2002 | Muscle | Yes | 17.342 | 0.089 |
| Yukon Aishihik 27602 | Northern mountain | Woodland | 2002 | Muscle | Yes | 15.873 | 0.079 |
| Yukon/Alaska Fortymile 27673 | Barren-ground | Grant's | 1994 | Muscle | No | 18.078 | -0.088 |
| Yukon Hart River 27703 | Northern mountain | Woodland | 2000 | Muscle | Yes | 16.126 | -0.052 |
| Yukon Hart River 27706 | Northern mountain | Woodland | 1999 | Muscle | Yes | 17.161 | -0.080 |
| Yukon Porcupine 27737 | Barren-ground | Grant's | 2001 | Muscle | No | 38.566 | -0.082 |
| Yukon Porcupine 27738 | Barren-ground | Grant's | 2001 | Muscle | No | 38.782 | -0.071 |
| Yukon Tay<br>27772 | Northern<br>mountain | Woodland | 2005 | Muscle | Yes | 18.863 | -0.025 |
| Yukon Tay<br>27773 | Northern<br>mountain | Woodland | 2002 | Muscle | No | 23.117 | -0.037 |
| British Columbia<br>Muskwa<br>28320 | Northern<br>mountain | Woodland | 2006 | Hide | Yes | 25.875 | -0.020 |
| British Columbia<br>Frog<br>28327 | Northern<br>mountain | Woodland | 2002 | Hide | No | 34.592 | -0.038 |
| British Columbia<br>Pink Mountain<br>28330 | Northern<br>mountain | Woodland | 2004 | Hide | Yes | 15.050 | -0.022 |
| British Columbia<br>Spatzizi<br>28332 | Northern<br>mountain | Woodland | 2003 | Hide | Yes | 22.504 | -0.032 |
| British Columbia<br>Chase<br>28336 | Northern<br>mountain | Woodland | 2004 | Hide | Yes | 19.111 | -0.017 |
| British Columbia<br>Frog<br>28337 | Northern<br>mountain | Woodland | 2003 | Hide | No | 36.985 | -0.008 |
| British Columbia<br>Tsenaglude<br>28348 | Northern<br>mountain | Woodland | 2006 | Hide | Yes | 23.783 | -0.028 |
| British Columbia<br>Itcha-Ilgachuz<br>28395 | Northern<br>mountain | Woodland | 2006 | Hide | No | 31.978 | 0.171 |
| British Columbia<br>Itcha-Ilgachuz<br>28402 | Northern<br>mountain | Woodland | 2006 | Hide | No | 34.767 | 0.151 |
| British Columbia<br>Atlin<br>28575 | Northern<br>mountain | Woodland | 2006 | Hide | No | 35.065 | -0.028 |
| British Columbia<br>Atlin<br>28580 | Northern<br>mountain | Woodland | 2006 | Hide | No | 35.118 | -0.013 |
| British Columbia<br>Columbia North<br>28646 | Southern<br>mountain | Woodland | 2014 | Hide | No | 35.525 | 0.045 |
| British Columbia<br>Columbia North<br>28649 | Southern<br>mountain | Woodland | 2014 | Hide | No | 35.580 | 0.040 |
| Northwest<br>Territories Sahtú<br>region<br>35082 | Boreal | Woodland | 2015 | Muscle | No | 38.251 | -0.011 |
| Alberta A La<br>Peche<br>40092 | Central<br>mountain | Woodland | 2018 | Faecal | No | 10.286 | -0.055 |

Laboratory protocols for the mitochondrial DNA control region sequencing analysis (mtDNA control region) are described in detail in Klütsch et al. (2012; 2016). 1,832 samples were included for the mtDNA analysis (Figure 2; Table 2).

**FIGURE 2.**
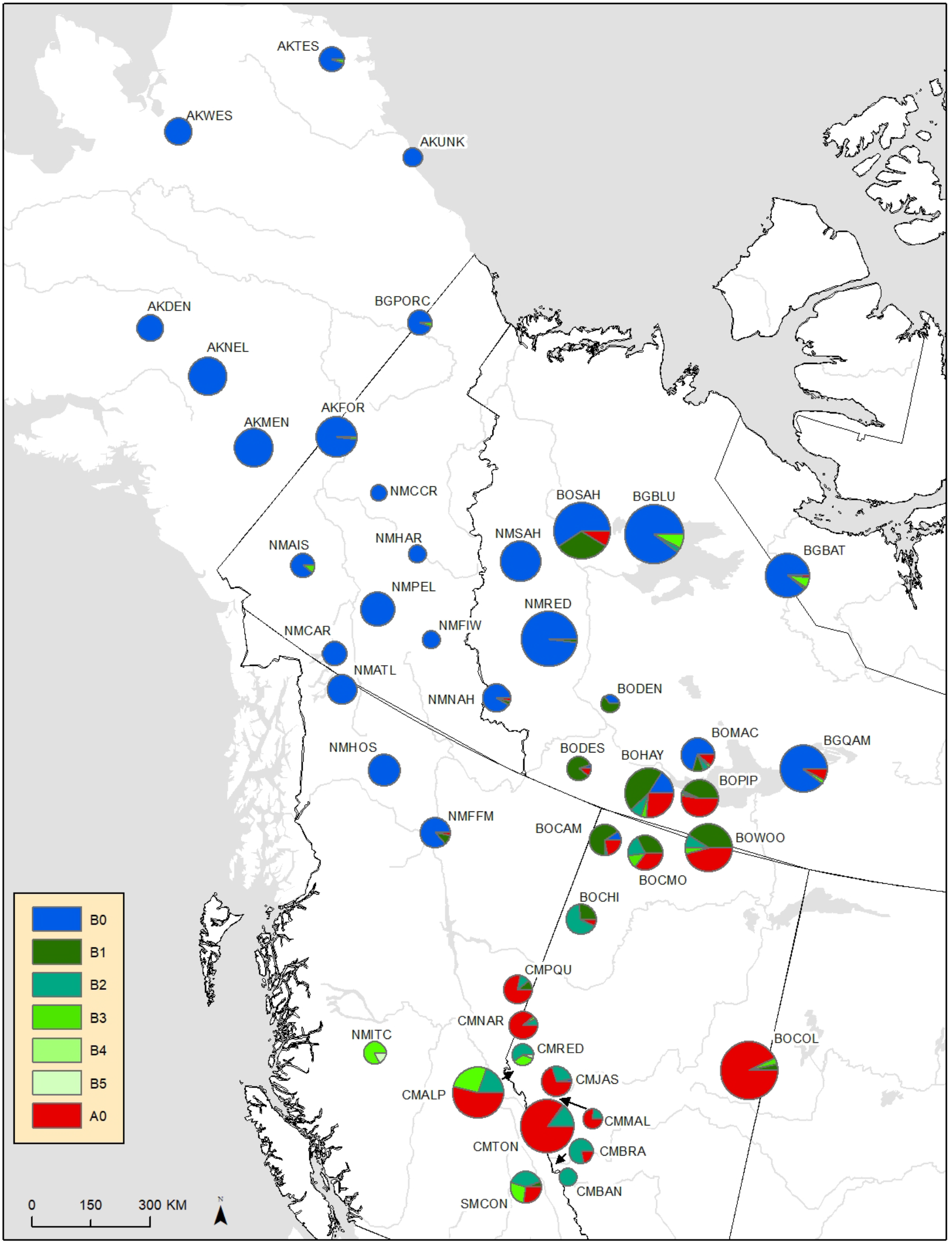
Map with the sampling locations for mitochondrial DNA analysis, showing the distribution of the mitochondrial control region haplotypes, indicating where haplotypes have introgressed (see Table 1 for locations and sample sizes).

**TABLE 2.**
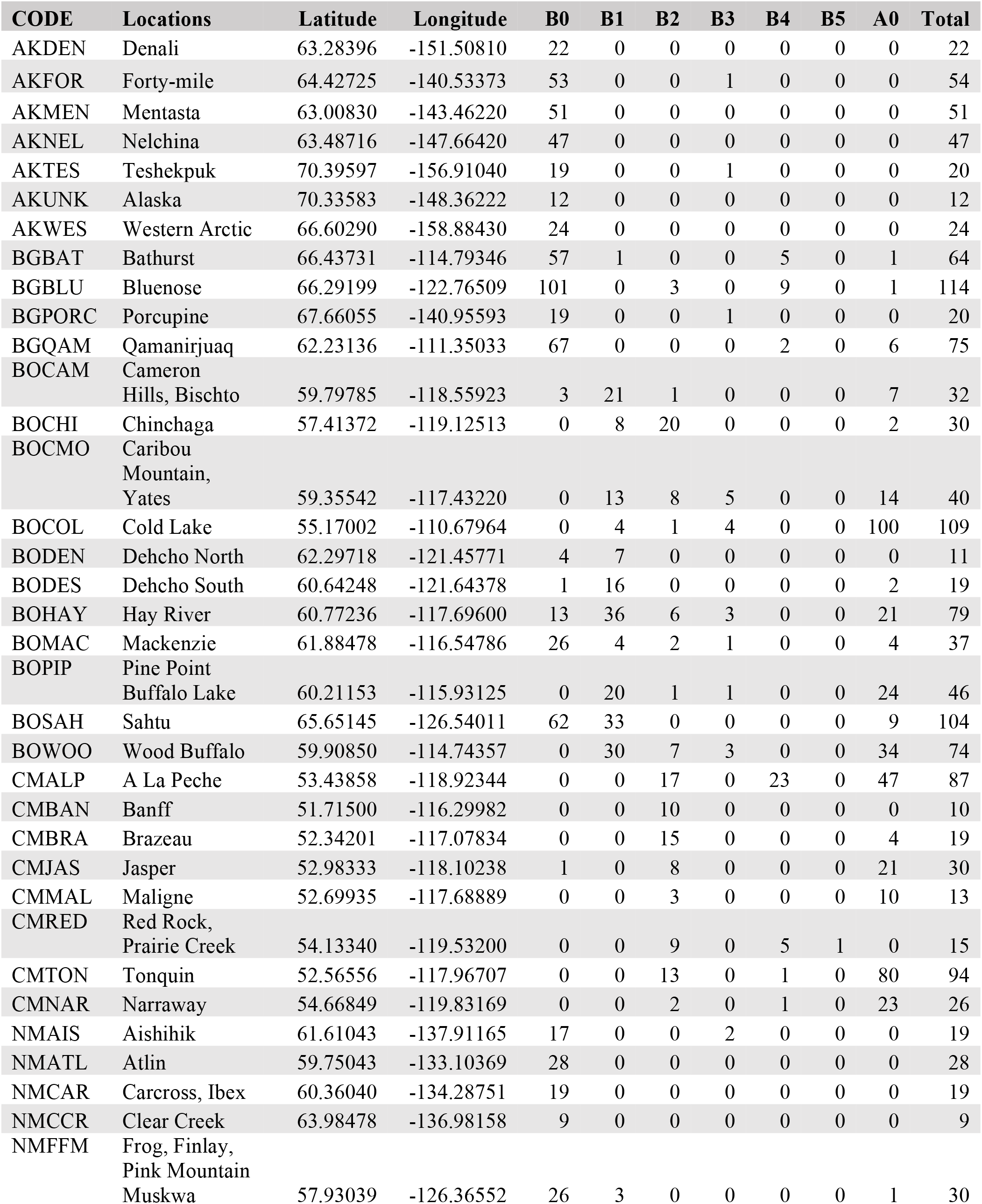
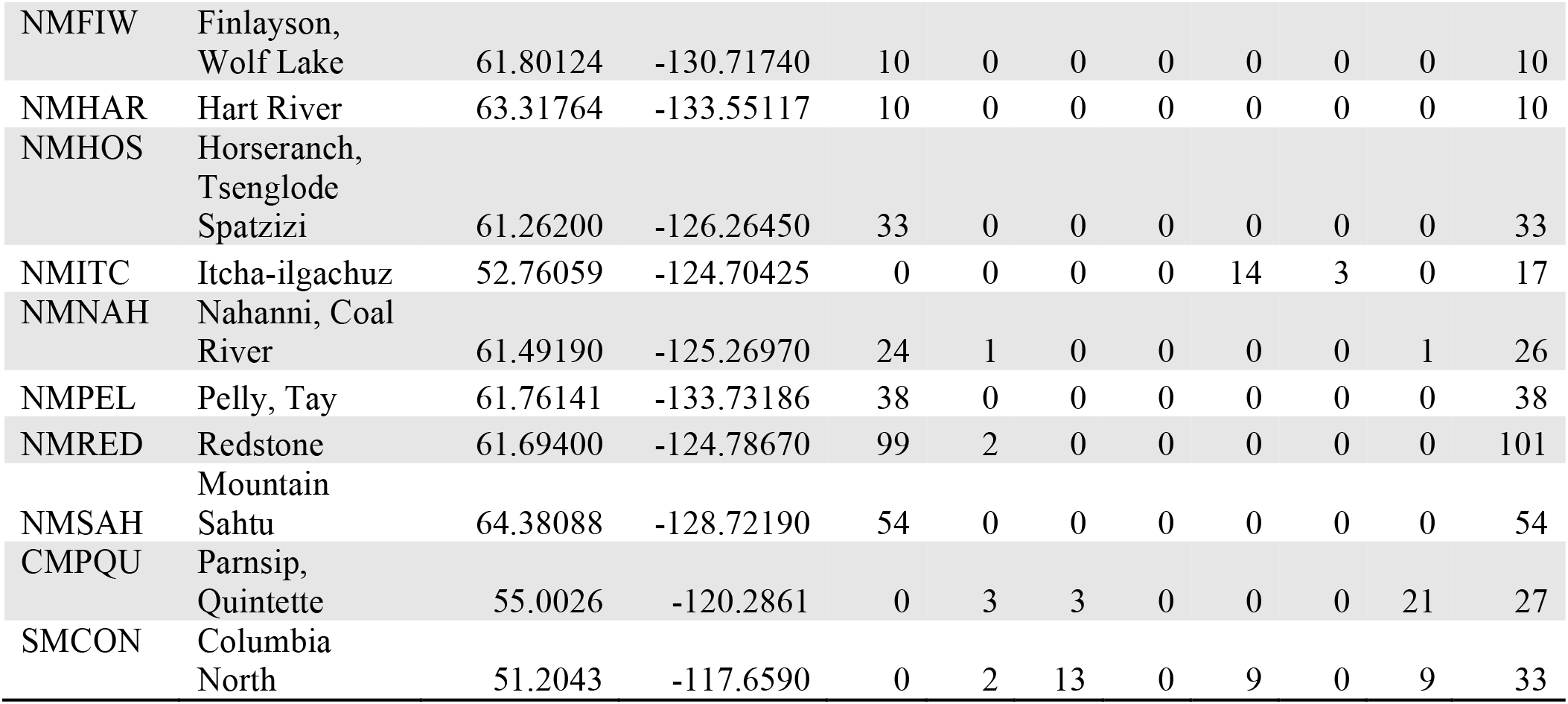
Mitochondrial control region sampling locations with their letter codes from Figure 2. Codes beginning with AK indicate samples from Alaska, BO indicates boreal caribou, CM central mountain, NM northern mountain, and SM southern mountain caribou. The number of each haplotype groups from each location is listed, as well as the total number of samples.

### 2.2| Mitochondrial DNA analysis

We assessed phylogenetic relationships and divergence times among mitogenome control region haplotypes using Bayesian methods. The software jModelTest 0.1.1 (Posada, 2008) was applied to identify HKY+G as the best substitution model using the Bayesian Information Criterion for caribou haplotypes. Two maximum clade credibility trees were created using BEAST v1.10.4 (Suchard et al., 2018) using time calibrated tips from ancient DNA derived haplotypes under a strict clock model, HKY+G substitution model, default optimization schedule, MCMC chain-length of 200 million, sampling every 20,000 generations and removing the first 10% of runs. The two independent runs were combined using the BEAST v1.10.4 package LogCombiner. We analyzed results from BEAST in Tracer v1.7 (Rambaut, Drummond, Xie, Baele, & Suchard, 2018) and all effective sample sizes (ESS) were much greater than 200, indicating length of MCMC in accurately representing the posterior distribution was appropriate (Kuhner, 2009). The phylogenetic trees we estimated were summarized in the BEAST v1.10.4 package TreeAnnotator and visualized in FigTree 1.4.4 (Availabale: http://tree.bio.ed.ac.uk/software/figtree/). Divergence times were calculated as the node heights of the 95% highest posterior density (HPD) intervals.

### 2.3| Genome filtering and variant calling

We used Trimmomatic version 0.38 (Bolger, Lohse, & Usadel, 2014) to trim adaptors and other Illumina sequences from the reads using the sliding window approach (4 base pairs at a time) to trim reads once the phred score dropped below 15. We aligned the filtered reads to the reference genome (Taylor et al., 2019) using Bowtie2 version 2.3.0 (Langmead & Salzberg, 2012). The resulting SAM files were then converted to BAM files and sorted using Samtools version 1.5 (Li et al., 2009). We removed duplicate reads and added correct read group information to each BAM file using Picard version 2.17.3 (Available: http://broadinstitute.github.io/picard/) and then re-sorted the BAM file in Samtools and built an index using Picard.

We called variants using Haplotype Caller in GATK version 3.8 (McKenna et al., 2010). This produced a variant call format (VCF) file for each caribou. These were combined, including with the genomes from Taylor et al. (2020), using the Combine GVCFs function. We performed joint genotyping using Genotype GVCFs also in GATK. To ensure only high quality variant sites were included in subsequent analyses, we did two rounds of filtering in VCFtools version 0.1.14 (Danecek et al., 2011). First, we removed indels and any site with a depth of less than 5 or more than 80 (double the depth we were aiming for across the genome for the genomes from Taylor et al., 2020), and removed any low-quality genotype calls (minGQ) and low quality sites (minQ), with scores below 20 (these are changed to missing data in VCFtools). Second, we filtered to remove all missing data. The resultant VCF file contained 7,486,778 single nucleotide polymorphisms (SNPs). We measured the mean depth for each individual in VCFtools. To estimate individual inbreeding coefficients, we pruned the data in Plink (Purcell et al., 2007) to remove SNPs with a R^2^ value of greater than 0.1 with any other SNP within a 50 SNP sliding window, advancing by 10 SNPs at a time, in order to minimize linkage. We used Plink to calculate the inbreeding coefficients (--het command).

We also produced a VCF file containing a Sitka deer genome (*Odocoileus hemionus sitkensis*) to use as an outgroup in some analyses. The raw reads for the Sitka deer, which was sequenced as part of the CanSeq150 Initiative, were downloaded from the NCBI database (Bioproject PRJNA476345, run SRR7407804). The reads were aligned to the reference genome and filtered as with the other genomes. We used Combine GVCFs and then performed joint genotyping using Genotype GVCFs in GATK to produce a VCF file with all caribou and the Sitka deer which was filtered in VCFtools as before. The resultant VCF file contained 8,048,723 SNPs.

### 2.4| Genome demographic and phylogenomic analyses

We used Pairwise Sequentially Markovian Coalescent (PSMC; Li & Durbin, 2011) to reconstruct historical changes in population size. Using the BAM files, we made a consensus fastq file for each caribou using the Samtools and BCFtools 1.5, filtering to remove sites with a depth of less than 10 or more than 80. We did not include the lowest coverage individual from Cold Lake as PSMC is particularly sensitive to coverage (Nadachowska-Brzyska, Burri, Smeds, & Ellegren, 2016). The consensus fastq was converted into an input file for PSMC (using the ‘fq2psmcfa’ command), adding a filter to remove anything with a mapping quality score below 20. Following advice from the PSMC manual (https://github.com/lh3/psmc) and Nadachowska-Brzyska et al. (2015), we used several pilot runs to optimize the input parameters (-t, -p, and -r), making sure that at least 10 recombination events were inferred at each interval after 20 rounds of iterations. We found that the free atomic time intervals (-p) and the initial value of r = θ/ρ (-r) did not change the performance or the results. However, we found that the program performed better when setting the upper limit to the time to most recent common ancestor (-t) to 5. We then performed 100 bootstraps for each individual to estimate the variance in the inferred effective population sizes. We plotted the results using the mutation rate as calculated for reindeer of 1.1E-8 (Chen et al., 2019) and a generation time of 7 years (COSEWIC, 2014-2017).

Using the VCF file we performed a principal component analysis in R 3.4.4 (R Development Core Team, 2018) using the packages vcfR (Knaus & Grünwald, 2017) and Adegenet (Jombart, 2008). We also ran subsets of individuals for higher resolution of some clusters (see Results) using VCF files made as above containing only those individuals.

For phylogenomic analyses, we generated both a consensus phylogeny reconstructed from conserved mammalian genes and a SNP phylogeny. For the former, we used BUSCO (Benchmarking Universal Single-Copy Orthologs 3.0.2; Waterhouse et al., 2018) to extract 4,104 conserved mammalian genes from the BAM files of each genome. Our reference genome reconstructed 3,820 (93.1%; Taylor et al., 2019) of these genes and so this was the upper limit for our re-sequenced caribou. The same was done for the Sitka deer and this was used as an outgroup to root the phylogeny. We used MUSCLE (Edgar, 2004) to align the sequences for each gene into fasta files, which were all run through RAxML 8 (Stamatakis, 2014) using the GTRGAMMA model and 1,000 bootstrap replicates. We then used ASTRAL-III (Zhang, Rabiee, Sayyari, & Mirarab, 2018) to create a consensus phylogeny. We used FigTree to visualize the consensus tree.

For the SNP phylogeny, we did not include the Sitka outgroup because it pushed the caribou too closely together to resolve the branching patterns, rather we rooted it where indicated by the BUSCO analysis. We used VCFkit (available here: https://vcf-kit.readthedocs.io/en/latest/, using numpy 1.14 as the programme does not work with newer versions) and generated a fasta file from the VCF file using the ‘phylo fasta’ command. The program concatenates SNPs for each sample, using the first genotype of each allele and replacing missing values with an N. We input the fasta file into RAxML and ran using the same parameters as above, visualizing the best tree in FigTree.

### 2.5| Genomic tests for introgression

We used Treemix 1.13 (Pickrell & Pritchard, 2012) to reconstruct a maximum likelihood phylogeny and to visualize migration events between populations. To create the input file from the VCF file we used Stacks 2.4.1 (Catchen, Hohenlohe, Bassham, Amores, & Cresko, 2013) using the populations function. We input this file into Treemix and ran 10 iterations of 0-9 migration events. To account for possible linkage we grouped the SNPs into windows. We plotted the resulting trees and residual plots in RStudio 1.0.136 (RStudio Team, 2015). We used the R package OptM (available here: https://cran.r-project.org/web/packages/OptM/index.html) to calculate the ad hoc statistic delta M, which is the second order rate of change in the log-likelihood of the different migration events, to help infer how many migration events to visualize. Because the OptM package requires different likelihood scores between iterations, we used two different window sizes in our runs, doing five iterations grouping SNPs in groups of 250 and five iterations grouping into groups of 500.

## 3| RESULTS

### 3.1| Mitochondrial phylogeny and signatures of introgression

Ancient mtDNA haplotype dates, indicated by the timing of tip termination, were included in the Bayesian phylogenetic reconstruction to infer divergence times of contemporary haplotypes. Importantly, BEL mtDNA associated with north-western North America coalesced approximately 120-130 kya (Figure 3). The overall mtDNA diversification is consistent with high effective population sizes given the number of emerging lineages over time and contemporary number of haplotypes. It is important to note that the haplotypes in this analysis are a subset of a much larger dataset that contains other regions of Canada as well as Eurasia, reflecting even more diversity.

**FIGURE 3.**
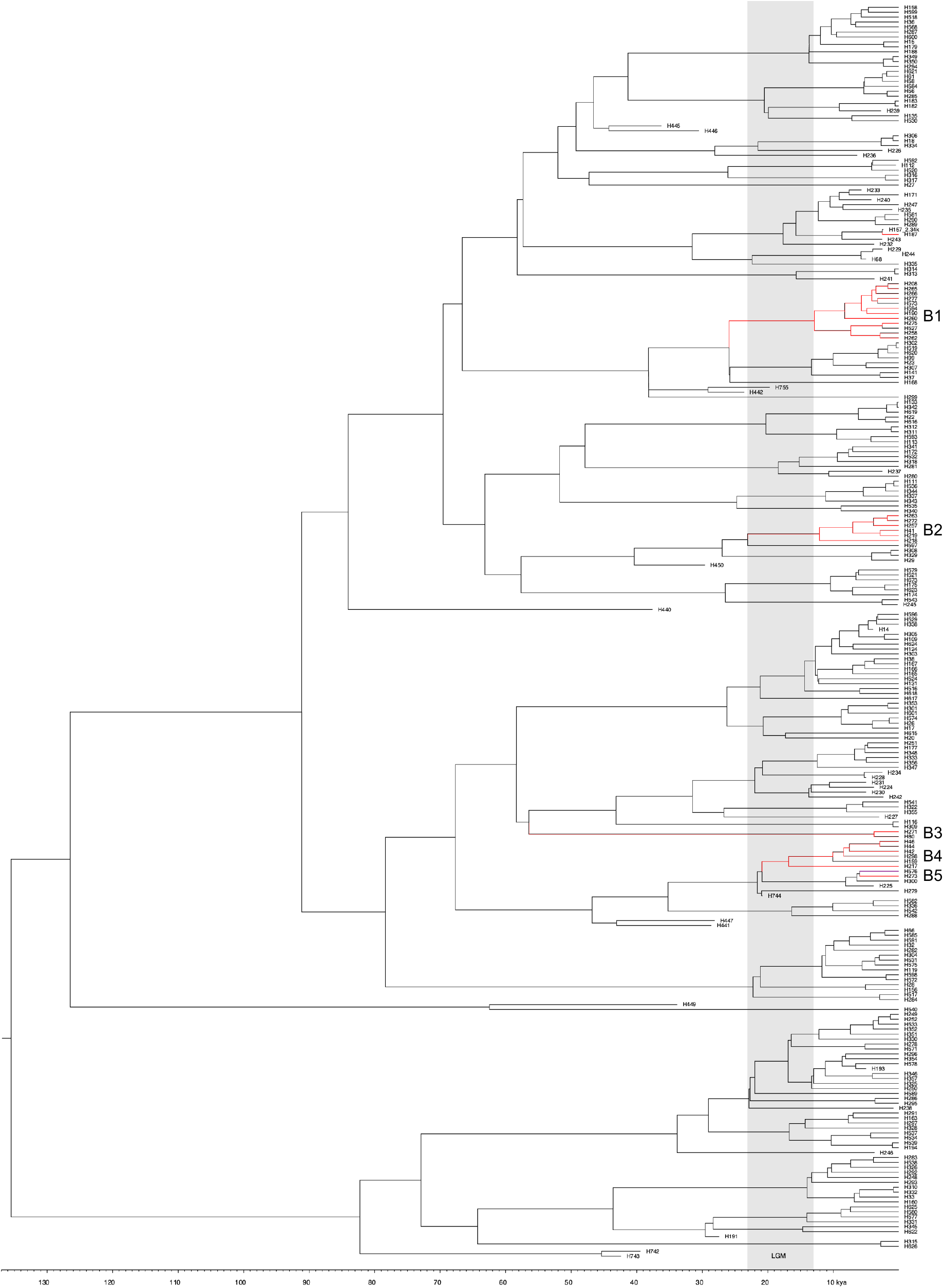
Mitochondrial haplotype phylogeny with the LGM shaded in gray. Introgression events are shown in red and refer to haplotypes from populations not from the same lineage.

Lineages A0 and B0 reflect the NAL and BEL, respectively (Figure 2). Lineages or haplotypes putatively reflecting introgression were identified based on clustering together and a predominant distribution within ecotypes not associated with Beringia (e.g. boreal caribou and southern mountain ecotypes; and a southern distribution such as Itcha-Ilgachuz or Jasper). Lineage B1 (Haplotypes 190, 208, 260, 265, 266, 277, 275, 527, 258, 262,573 and 594) dates back to approximately 38,000 to 25,000 years ago predating the LGM and reflecting more ancient introgression. Lineage B2 (Haplotypes 41, 218, 219, 247, 263 and 272) are associated with a broad distribution of boreal caribou in the west and into regions in Ontario. This lineage dates back to before the early LGM boundary. Lineage B3 (Haplotypes 80 and 271) associated with Caribou Mountain, Cold Lake, and Wood Buffalo boreal caribou are also found in Beringian regions such as the Fortymile and the Porcupine caribou populations. These haplotypes may reflect an ancient introgression event dating as far back as 60,000 years ago, significantly predating the LGM. Lineage B4 (Haplotypes 42, 44, 46, 159, 217, 298), associated with southern and central mountain caribou, dates to within the LGM and may reflect introgression that occurred from ancestral BEL haplotypes that moved south prior to the maximal ice sheet, post-LGM cannot be ruled out, however. The proliferation of mountain specific haplotypes in this lineage does support a more ancient introgression. Lineage B5 (Haplotypes 273 and 576) is associated with Itcha-Ilgachuz and Redrock-Prairie Creek is related to Lineage B4 and may reflect post-LGM introgression (Figure 2 and 3).

### 3.2| Genomic structure and demographic history

Our 33 whole genome sequences were generally recovered to high depth, with the average depth for SNPs in the VCF file ranging between 8.4 and 40.1 X coverage (Table 1). The lowest coverage individual, a boreal caribou from Cold Lake, was one of the three reconstructed from faecal DNA, and the results for this individual for some analyses may need to be interpreted with caution. The inbreeding coefficients varied between individuals, with most caribou having very low values (Table 1). Elevated coefficients were seen in the northern mountain Itcha-Ilgachuz population (0.171 and 0.51) and slightly elevated in the Aishihik population (0.089 and 0.079; Table 1). The coefficient was lower for the lowest coverage Cold Lake genome (−0.260), as compared to the Cold Lake individual with better coverage (0.012), which may indicate some error for this individual (Table 1).

Demographic reconstruction showed expansion and differential historical population trajectories starting at ~120 kya (Figures 4a-h; see Figures S1-4 for bootstrap plots). For boreal caribou, the individuals from the Northwest Territories had a much higher historical population size during the expansion than the individual from Cold Lake (Figure 4a and Figure S1 a-c). This agrees with previous results indicating parallel evolution of the BEL boreal caribou from the Northwest Territories and the NAL boreal caribou from more southern parts of the range (Polfus et al., 2017; Taylor et al., 2020).

**FIGURE 4.**
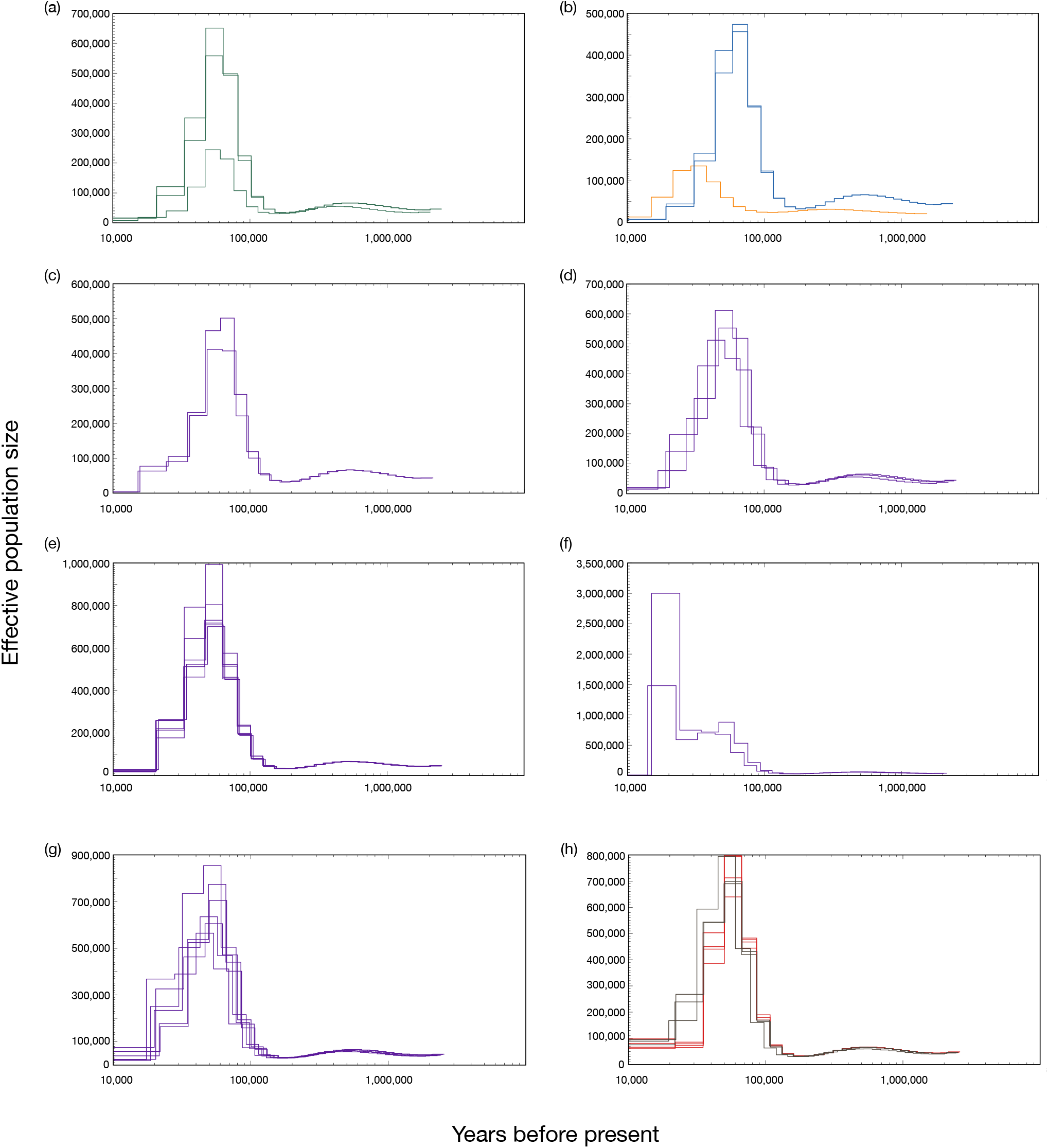
PSMC reconstruction of historical population sizes of (a) boreal caribou, with the Cold Lake caribou showing much lower peak effective population sizes than those from the Northwest Territories, (b) central (orange) and southern mountain (blue) caribou, (c) Northern mountain Itcha, (d) Northern mountain Chase, Pink Mountain, and Muskwa, (e) Northern Mountain Frog, Atlin, Spatzizi, Tseneglode, (f) Northern Mountain Aishihik, (g) Northern Moutain Tay, Redstone, Hart River, (h) Barren-ground (red) and Grant’s (brown). Note differences on the Y axis showing effective population sizes.

Caution may be needed with the interpretation of some individuals given that both coverage and high levels of inbreeding can impact PSMC results (Nadachowska-Brzyska et al., 2016; Mather, Traves, & Ho, 2020), however, most individuals had coverage close to or above the 18X recommended (Table 1; Nadachowska-Brzyska et al., 2016). Itcha-Ilgachuz and Aishihik had elevated inbreeding coefficients which may impact results, and we did see large error margin predicted by the bootstrap analysis for Aishihik individuals, which have a very different demographic reconstruction than the others (Figure 4f and Figures S3a and b). However, removing runs of homozygosity did not change PSMC reconstructions in avian species (Nadachowska-Brzyska et al., 2015; Nadachowska-Brzyska et al., 2016), and both Aishihik individuals gave the same shape of curve.

The central mountain caribou had the lowest overall population sizes during expansion with a maximum peak around 140,000 (Figure 4b and Figure S1d), but with lower coverage across the genome (~10X) perhaps impacting the result. Southern mountain caribou reached around 450,000 individuals (Figure 4b and Figure S1 e-f). The historical effective population sizes varied widely within the northern mountain caribou. Itcha-Igachuz and Pink Mountain had the lowest peak population sizes, similar to the southern mountain caribou (Figure 4c-d and Figure S2 a-c). The other northern mountain caribou had peak effective population sizes ranging from ~500,000 to 1 million individuals (Figures 4d-g, Figure S2 d-k, Figure S3 a-h), with the exception of Aishihik with indicated population sizes peaking at 1.5 and 3 million caribou (Figure 4f), although with a large variation observed around these values in the bootstrap analysis (Figure S3 a-b). The Aishihik herd is near the north-western edge of the northern mountain range (Figure 1), and further sequencing in the region would be beneficial to clarify the unique demographic history uncovered pointing to a later population expansion, occurring during the LGM (Figure 4f and Figures S3a and b).

Barren-ground and Grant’s caribou (which both belong to the barren-ground DU) had similar historical population trajectories, with peak population sizes between 650,000 and 800,000 individuals (Figure 4h and Figure S4 a-g). Population sizes reached their highest at around 50-60 kya for boreal caribou (Figure 4a), 50-80 kya for southern mountain (Figure 4b), typically around 50-80 kya for northern mountain caribou (Figures 4c-g), apart from Aishihik with occurred much later at around 15-25 kya (Figure 4f), and around 50-70 kya for barren-ground and Grant’s caribou (Figure 4h). All populations had large declines in population sizes starting between 30-50 kya, aside from Aishihik for which population size declines occurred around 15 kya

The PCA with all individuals separated the boreal Cold Lake individuals and the northern mountain caribou from Itcha-Ilgachuz most strongly (Figure 5). Southern and central mountain caribou formed separate clusters, as did the boreal caribou from the Northwest Territories and northern mountain Pink Mountain caribou. The barren-ground caribou and the northern mountain Aishihik both formed clusters separating from the large cluster with all other northern mountain individuals and the Grant’s caribou (Figure 5). The PCA with only the northern mountain caribou strongly separated Itcha-Ilgachuz and Aishihik (Figure 6a) which may be related to their elevated inbreeding coefficients (Table 1). When removing these individuals, we see four major clusters: 1) Atlin, 2) Pink Mountain, 3) the other herds from British Columbia (Chase, Spatzizi, Frog, Muskwa, and Tseneglode), and 4) the herds from the Yukon and the Northwest Territories (Tay, Hart River, and Redstone; Figure 6b).

**FIGURE 5.**
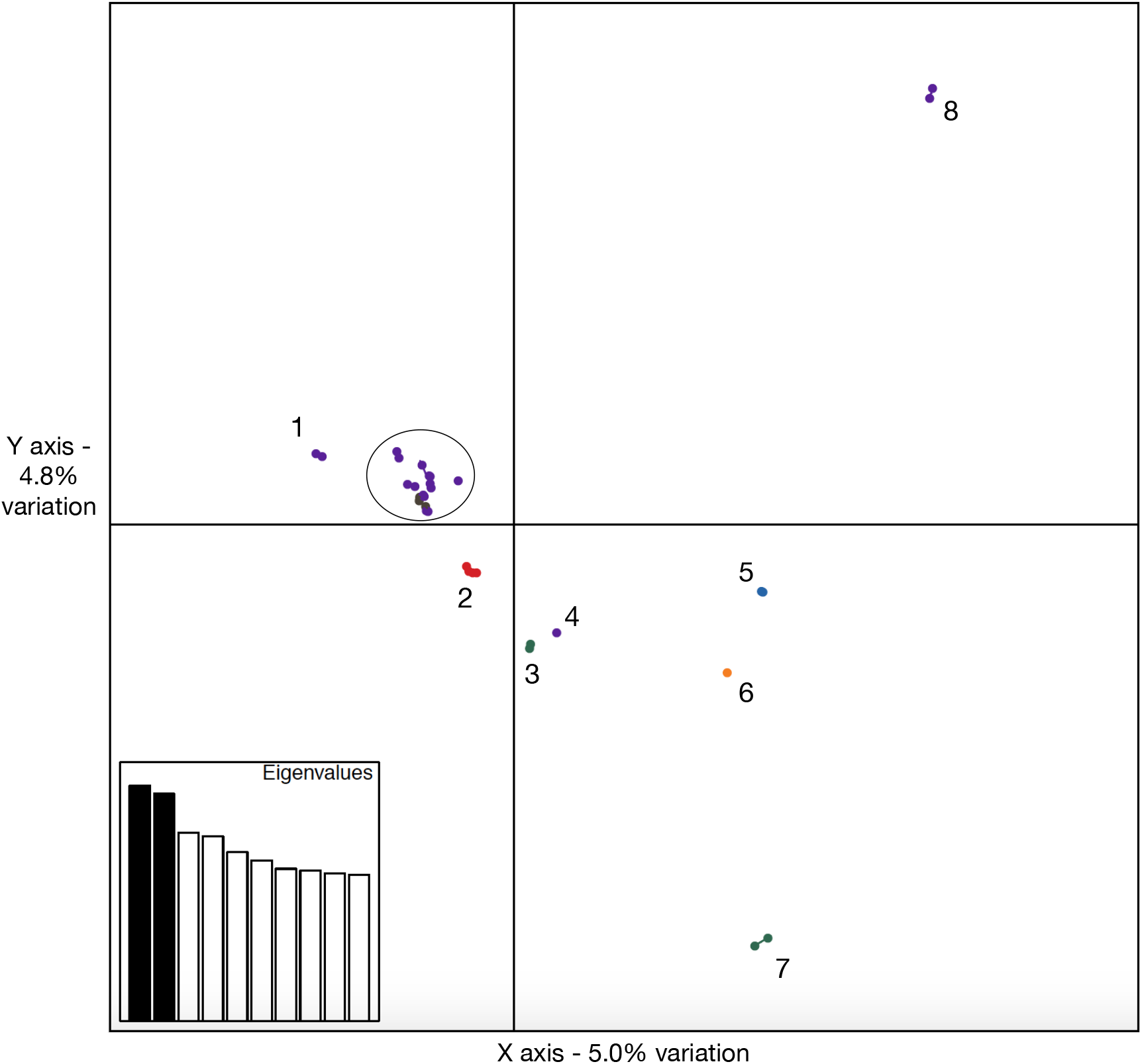
PCA plot with all caribou genomes. 1. Northern mountain Aishihik; 2. All four barren-ground caribou; 3. Boreal caribou from the Northwest Territories; 4. Northern mountain Pink Mountain; 5. Southern mountain Columbia North; 6. Central mountain A La Peche; 7. Boreal Cold Lake; 8. Northern mountain Itcha-Ilgachuz. All other northern mountain and the Grant’s caribou are within the circle.

**FIGURE 6.**
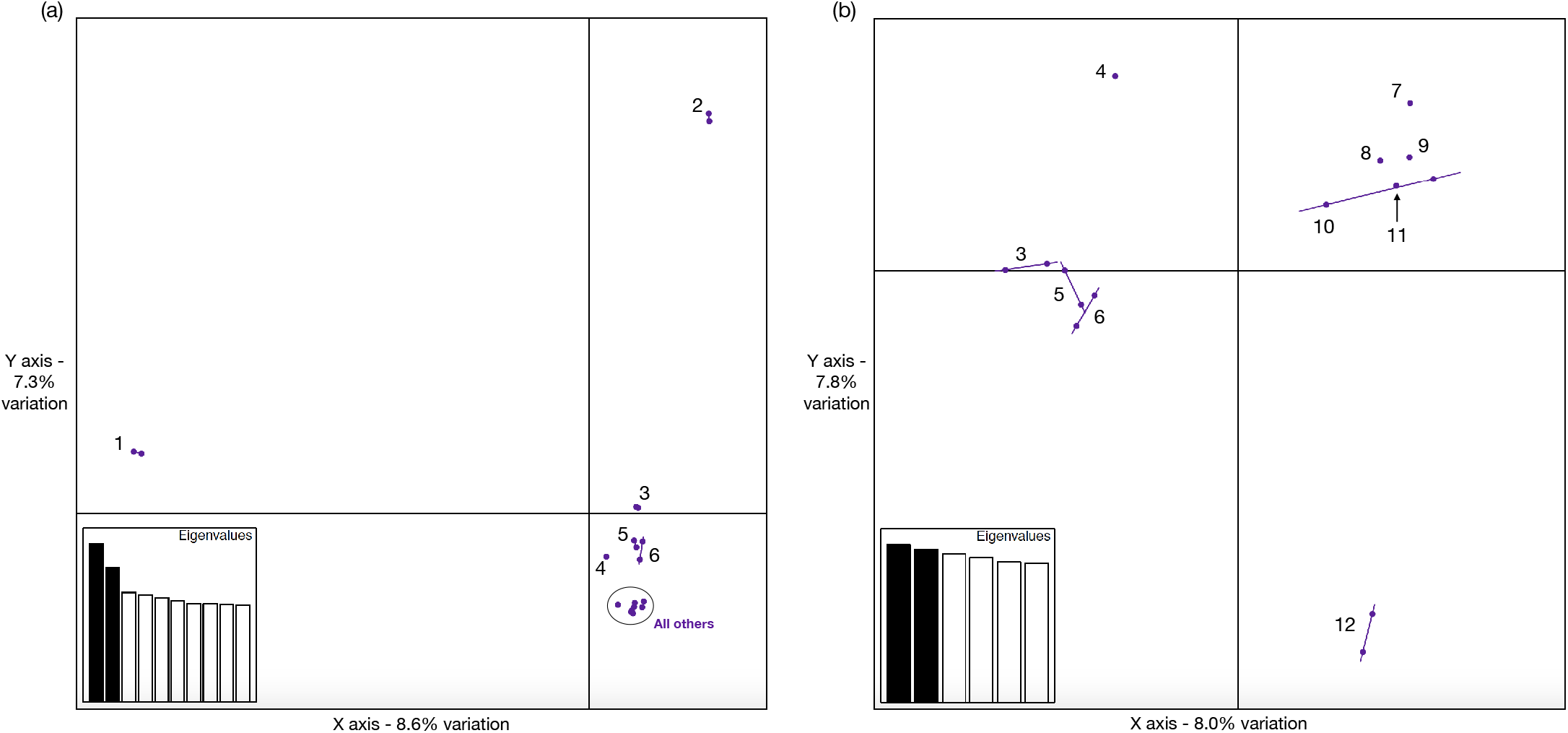
PCA plot showing only the northern mountain caribou (a) and showing the northern mountain caribou without Itcha-Ilgachuz and Aishihik individuals (b). 1. Itcha-Ilgachuz; 2. Aishihik; 3. Hart River; 4. Pink Mountain; 5. Redstone Northwest Territories; 6. Tay; 7. Chase; 8. Muskwa; 9. Spatzizi; 10. Frog; 11. Tseneglode; 12. Atlin.

The rooted phylogenomic reconstruction using conserved mammalian genes split the boreal caribou from Cold Lake as a separate clade from all others (Figure S5). The relationships within these two major clades have low support, however, which is unsurprising given it is built using conserved gene sequences. For finer scale resolution, we produced a phylogeny from the SNP data and rooted the tree where indicated by the conserved gene tree (Figure 7). Most of the northern mountain caribou group as sister populations although with the Grant’s caribou within them. All the northern mountain caribou from B.C., apart from Pink Mountain and Itcha-Ilgachuz, sit together in a clade, which is recovered as sister to northern mountain caribou from Tay in the Yukon and those from the Northwest Territories (which are neighboring herds geographically; Figure 1). Aishihik and Hart River northern mountain caribou sit with the Grant’s caribou, which matches the geography as these are the closest to the Grant’s caribou herds (Figures 1 and 7). The barren-ground caribou sit outside of the northern mountain and Grant’s caribou. Southern mountain caribou sit with the northern mountain caribou from Itcha-Ilgachuz, with boreal caribou from the Northwest Territories as a basal group, followed by the northern mountain caribou from the Pink Mountain herd, and then the central mountain caribou (Figure 7), patterns which do not closely match the geography (Figure 1) nor the DU designations.

**FIGURE 7.**
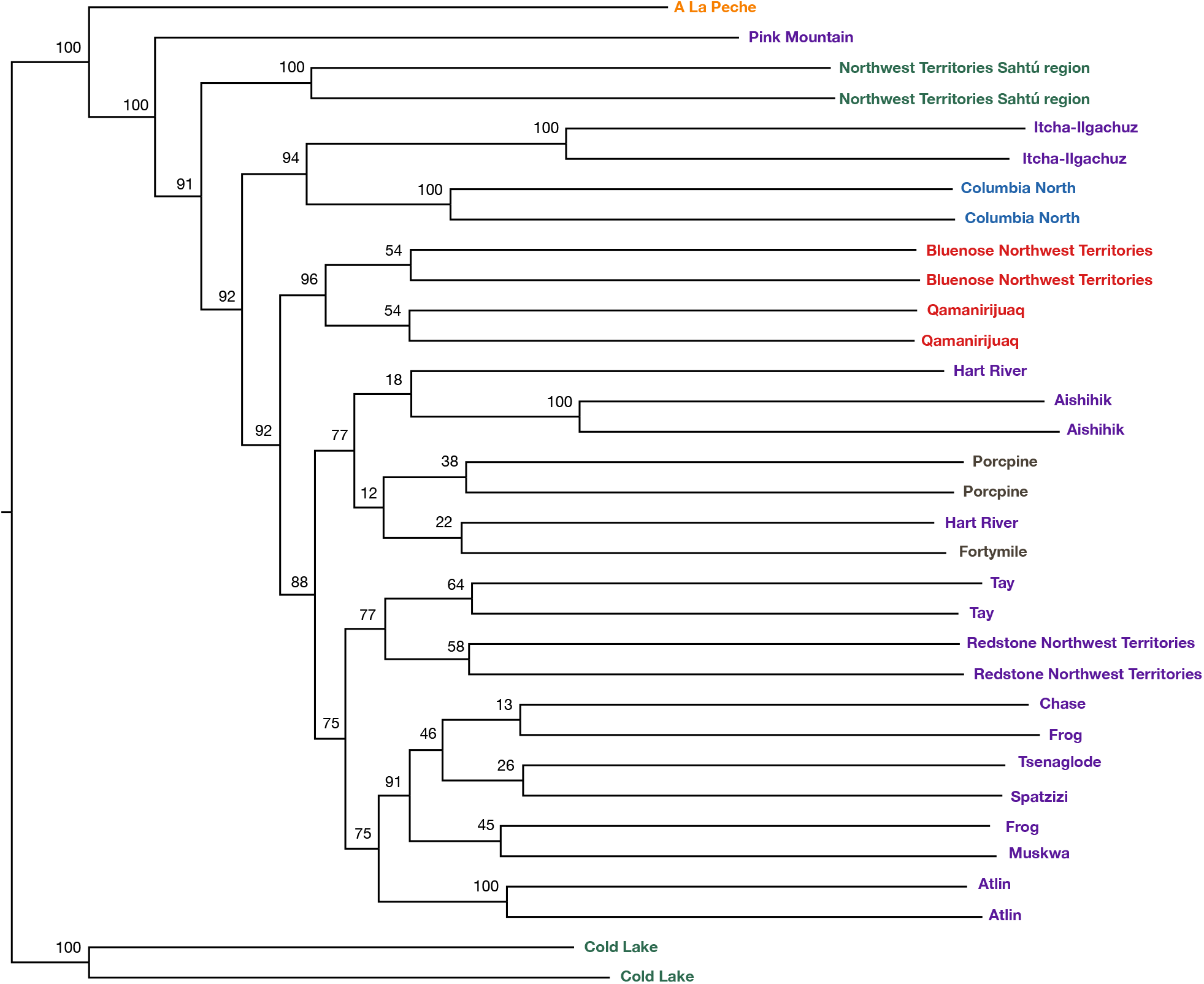
Maximum likelihood phylogenomic reconstruction from SNP data. The tree has been rooted where indicated by the BUSCO analysis, and bootstrap support values are indicated on the nodes.

### 3.3| Genomic signatures of introgression

To determine if some of the population and phylogenomic results could have been influenced by introgression, we first ran Treemix to infer migration events. Adding two migration events gave the highest delta M score (Figure S6), however when visualizing the trees inferring two migration events there was variation in where the events were inferred. Out of the 10 iterations, there were 14 different migration events inferred (out of a possible 20 with each iteration inferring two). The migration events that were inferred the most were from boreal Cold Lake into northern mountain Pink Mountain, and northern mountain Itcha-Ilgachuz into the ancestral population of central mountain A La Peche and southern mountain Columbia North (both inferred three times; all trees Figures S7a-j). Migration from southern mountain Columbia North into central mountain A La Peche, and from boreal Cold Lake into the ancestral population of central mountain A La Peche and southern mountain Columbia North were both inferred twice. The other 10 inferred instances all occurred once, with nearly all inferred migration events also occurring between populations not matching expected geographical patterns: boreal Cold Lake, boreal Northwest Territories, northern mountain Pink Mountain and Itcha-Ilgachuz, southern mountain Columbia North, and central mountain A La Peche (Figures S7a-j for full results), indicating that introgression likely impacted the phylogenomic reconstruction. The Treemix results also show a large drift parameter for northern mountain Aishihik and Itcha-Ilgachuz herds (Figures S7a-j), in line with the elevated inbreeding coefficients (Table 1).

## 4| DISCUSSION

### 4.1| Western caribou genetic structure and diversification

We used 33 whole-genome sequences and over 1,800 mitochondrial control region sequences to investigate genomic structure, patterns of introgression, and diversification in caribou from different regions of north-western North America. Overall genomic structure shows separation between individuals from different DUs, although with a few exceptions: boreal caribou from the Northwest Territories are not most closely related with the boreal caribou from Cold Lake (Figure 5 and 7), a reflection of parallel evolution of ecotype inferred previously (Horn et al., 2018; Polfus et al., 2017; Taylor et al., 2020). Northern mountain caribou have sub-structuring within them, notably Itcha-Ilgachuz and Aishihik are well separated in the PCA (Figure 5 and 6b). Pink Mountain caribou cluster closer to the Northwest Territories boreal caribou (Figure 5 and 7). The other northern mountain caribou follow a geographical pattern, with individuals from B.C. clustering together, with the exception of Atlin, which is found slightly to the west and north than the others, and individuals from the Yukon and the Northwest Territories, which are geographically closest also clustering together (Figure 6b). The populations not found where expected in the phylogenomic reconstruction, based on their geographic locations, were likely impacted by introgression events. For example, introgression from boreal caribou from Cold Lake into boreal caribou from the Northwest Territories, and into Pink Mountain, as well as introgression between Itcha-Ilgachuz and central and southern mountain caribou, was found in Treemix (Figure S7).

Genetic sub-structuring within the northern mountain ecotype is also apparent from the PSMC reconstruction, with different demographic histories being reconstructed (Figures 4c-g). This highlights the need to analyze multiple individuals of the same species, or even ecotype. Interpretations based on single individuals, while common with PSMC, can be biased (Kozma et al., 2016). Caribou show rising and differential effective population sizes starting at ~120 kya (Figure 4), consistent with the mtDNA coalescent timing (Figure 3). The overall patterns from the mitochondrial data, the PCA, nuclear phylogeny, and the PSMC analysis demonstrate that there is no one mountain ancestor, and that even geographically close populations may have different demographic histories. It is likely that mountain caribou underwent multiple different colonization events which influenced the current genetic structure and biogeography, given that there were multiple periods after the original diversification when ice-free corridors would have allowed the movement of these highly mobile animals (e.g. Figure 4b-c). Itcha-Ilgachuz caribou appear to be from a Beringian origin, with mostly B mitochondrial haplotypes, but may have undergone a more ancient colonisation, followed by introgression, reflecting their divergence from other northern mountain caribou. Southern mountain caribou may also be of Beringian origin, having mostly B haplotypes, and may also have undergone a more ancient colonisation. In contrast, central mountain caribou have more A haplotypes, and so are likely of NAL origin. The genetic groupings we see in the rest of the northern mountain caribou also likely demonstrate different histories, with a clade showing affinity with the Grant’s caribou, and a group with Redstone and Tay (Figures 5–7, also clear when viewing the third and fourth PCA axes with all individuals Figure S8). We also see a clade with the British Columbia northern mountain caribou herds. Although Atlin is positioned basally on the phylogeny, and they separate on the PCAs (Figures 5–7), it is possible they colonised North America separately from the others. It is clear that mountain caribou comprise populations which have been differentially impacted by potentially differential colonization timing with subsequent introgression, and are not one homogenous group.

As well as demonstrating different demographic histories, the PSMC results show that effective population sizes reached up to ~1 million individuals for some populations, (Figure 4, S1-4), indicating an incredible diversification of caribou in the north-western regions of North America long before the LGM, congruent with the large number of mtDNA haplotypes which emerge (Figure 3). The large historical effective population sizes are far greater than most PSMC reconstructions of mammal species thus far (e.g. Bunnefeld, Frantz, & Lohse, 2015; Ekblom et al., 2018; Liu et al., 2018; Mays et al., 2018; Miller et al., 2012; Tsuchiya, Dikow, & Cassin-Sackett, 2020; Westbury et al., 2018; Westbury, Petersen, Garde, Heide-Jørgensen, & Lorenzen, 2019; Yim et al., 2013; Yuan et al., 2018), with the potential exception of some bat species, depending on the generation time used (Chattopadhyay et al., 2019).

Especially for a large mammal species, these high peak effective population sizes likely had lasting impacts on contemporary diversity. Many caribou populations have very high levels of genetic diversity, as seen from the very low whole genome inbreeding coefficients for most individuals (Table 1) but also previous studies using microsatellites which found high levels of diversity for a large mammal species (e.g. Courtois, Bernatchez, Ouellet, & Breton, 2003; McLoughlin, Paetkau, Duda, & Boutin, 2004; Zittlau, Coffin, Farnell, Kuzyk, & Strobeck, 1998). In addition, the variation in morphology and life history (COSEWIC, 2011) may also have originated during the diversification event. Further sampling would be needed to determine how closely the pre-LGM population expansion is correlated with contemporary diversity.

It is possible that the PSMC peak is being inflated by population structuring (Mather et al., 2020), but even so, ~120-50 kya was clearly a critical period of diversification between lineages, as shown from both the nuclear and mitochondrial DNA. Hence, prevailing theory about the evolution of caribou lineages in refugia during the LGM, followed by recolonization and a ‘hybrid swarm’ in the mountains (McDevitt et al., 2009; Yannic et al., 2013) is likely oversimplified, further bolstered by the evident sub-structuring within the northern mountain ecotype (Figure 6a-b), and the evidence from the mtDNA of multiple introgression events occurring before, as well as after, the LGM (Figure 2 and 3).

### 4.2| Relationship of caribou demographic history to temperature and glaciation

Determining how populations have responded to large fluctuations in climate and varying environments throughout the Quaternary may help us to understand how they could respond under future climate change (Chattopadhyay et al., 2019; Kozma et al., 2016; Kozma et al., 2018; Louis et al., 2020; Yannic et al., 2013). Particularly at northern latitudes, there were large changes in available habitat and levels of glaciation (Chattopadhyay et al., 2019) which would have influenced caribou distributions and abundance. Comparing the key dates of caribou population expansion, decline, and introgression events to temperature and ice coverage at the same time periods (Figure 8; maps reconstructed from data from Batchelor et al., 2019) could lead to some insights. Just before caribou population expansion was the Eemian, a period of large climatic oscillations (Kozma et al., 2016; Miller et al., 2012). The beginning of the caribou diversification coincides with the transition to the last glacial period ~110 kya, when interglacial temperatures changed from very warm at the end of the Mid-Brunhes event to a period of cooling (see Figure 2 in Kozma et al., 2016; Miller et al., 2012). Ice coverage was not very extensive over north-western North America during this time (Figure 8a), and so there was likely an abundance of available habitat for caribou to disperse into. The continued period of cooling coincides with the peak of the PSMC plots during the diversification, usually around 50-80 kya (Figure 4; Kozma et al., 2016; Schmidt & Hertzberg, 2011).

**FIGURE 8.**
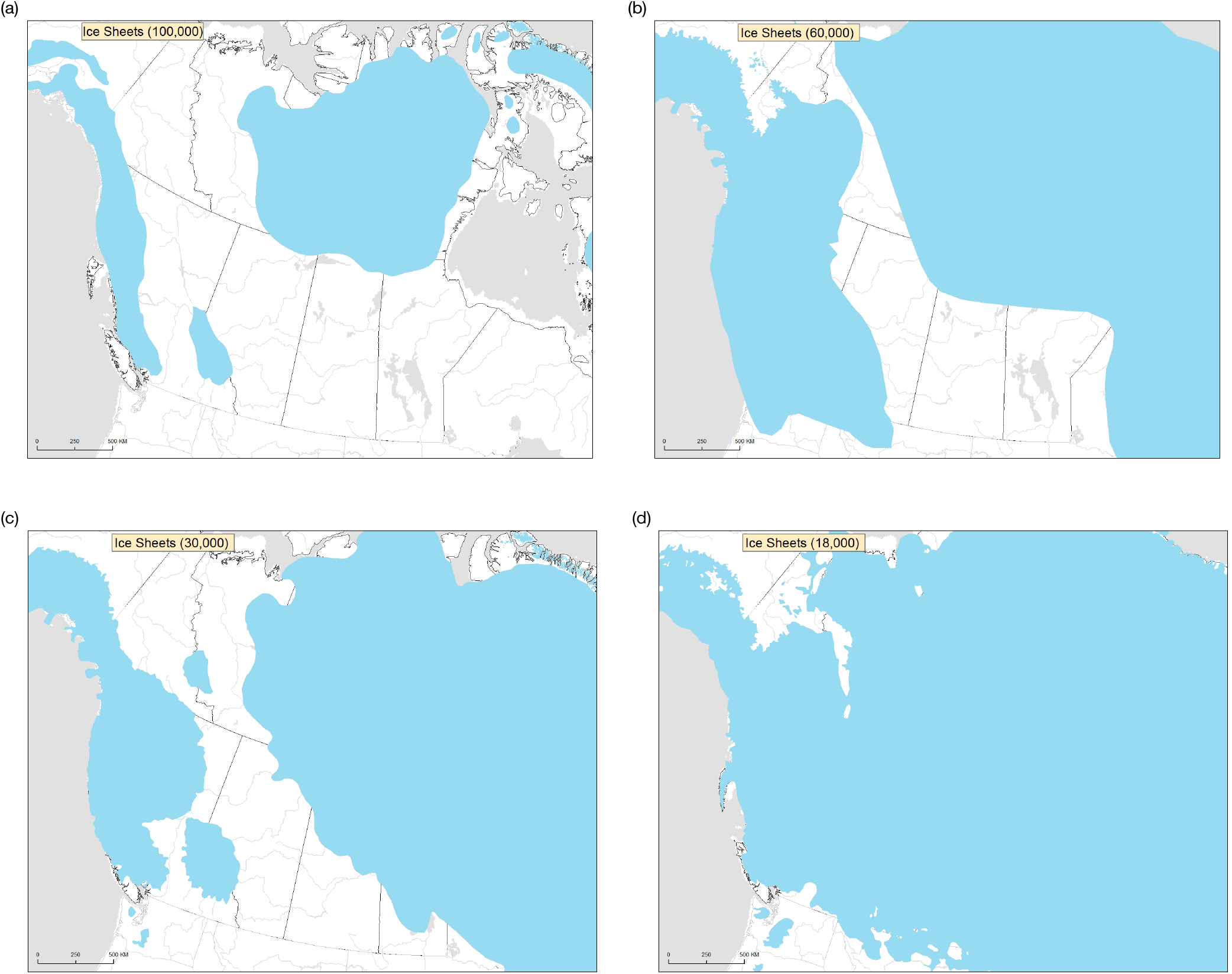
Maps showing the extent of Ice coverage in North America at 100 ky (a), 60 kya (b), 30 kya (c), and 18 kya (d), key time points of divergence and introgression events for caribou.

Interestingly, the dates of the pre-LGM introgression events inferred from the mtDNA, ~60 kya and ~30 kya (Figure 3), coincide with periods of lower temperatures during the temperature oscillations of the last glacial period (Kozma et al., 2016; Schmidt & Hertzberg, 2011) and increased ice coverage with an ice-free corridor (Figure 8b and c). Range re-distributions under more extensive glaciation may have increased the likelihood of introgression events among caribou populations normally geographically separated, even if population sizes were not necessarily small, as was the case at 60 kya (Figure 4). The ice-free corridor would have allowed movement, and channelled the movement along a specific path, perhaps facilitating introgression events between populations on either side of the ice sheets. Another two of the introgression events date to around the LGM, another period of maximal glaciation (Figure 8d).

The strong decline in effective population sizes generally coincides with periods of particularly rapid and dramatic climate changes of up to 16 °C, with interstadial warming events (known as Dansgaard-Oeschger, or D-O, cycles; Schmidt & Hertzberg, 2011; Cooper et al., 2015). During this period, from ~50 kya, North America lost around 72% of its large mammalian genera likely due to these rapid warming events (Cooper et al., 2015; Lorenzen et al., 2011). Human colonization did not occur in North America until 15 kya ruling out anthropogenic impacts (Lorenzen et al., 2011). It thus seems likely that the population declines we observe in caribou were also driven by these rapid changes in temperature and thus this species is likely vulnerable to future rapid climate warming.

Caribou effective population sizes were already much lower going into the LGM and remained stable after this period up until 10 kya (Figure 4) where PSMC loses accuracy. The barren-ground and Grant’s caribou maintained higher effective population sizes at this point (Figure 4h) never dropping as low as mountain and boreal caribou (the latter two both belong to the woodland subspecies). This suggests that caribou ecotypes which rely on forested habitat may have been more severely impacted by past climate changes, potentially due to being more closely tied to a habitat which cannot shift rapidly. Although historical population size changes were assumed to be due to climate changes, predators and other biotic interactions likely also play a role (Bai et al., 2018; Kozma et al., 2018). Woodland caribou may have declined further than barren-ground caribou due to differences in life history strategies, for example, the larger group sizes of barren-ground caribou could have buffered against changes in predator distributions. More recently, after the LGM caribou have again undergone population expansion and diversification, as evidenced by the number of new haplotypes emerging within the last 10 kya (Figure 3). This expansion coincides with a period of relatively stable temperatures (Schmidt & Hertzberg, 2011), and low glaciation which provided an abundance of habitat for caribou to diversify into, similar to the original diversification ~110 kya.

The Aishihik northern mountain caribou are an exception in that, in addition to the increase in population size beginning around 100 kya, they did not experience rapid declines and had a large increase in effective population size which appears to coincide with the LGM (Figure 4f). It is possible that they were in a separate region during the period of decline and only later colonised their current range. They may also have been in a different refugium from the other caribou sampled here during the LGM, as there were likely several sub-refugia within Beringia (Campbell et al., 2015; Galbreath et al., 2011; Scribner et al., 2003; Shafer et al., 2010). Further sampling will be needed to disentangle the unique demographic history of this population.

## 5| CONCLUSIONS

We have demonstrated, using nuclear whole-genome sequencing combined with large scale mitochondrial sequencing, that the major diversification of caribou occurred starting at the beginning of the last glacial period, ~110 kya. We also found genomic sub-structuring and evidence of multiple introgression events, indicating an evolutionary history more complex than diversification in two major refugia during the LGM, followed by more recent colonization and admixture (Klütsch et al., 2012; McDevitt et al., 2009; Weckworth et al., 2012; Yannic et al., 2013). Introgression events appear to be tied to the extent of glaciation and range re-distributions. Introgression events appeared to date to cold periods with high levels of glaciation often with ice-free corridors across North America. Population expansion occurred during a period of cooling with open habitat availability. Declines coincided with historical periods of temperature oscillations and rapid warming events, indicating that caribou may be vulnerable in future rapid climate warming. This may be especially true for woodland ecotypes which declined to lower effective population sizes than barren-ground caribou. Currently, woodland ecotypes continue to have smaller, in many cases declining population sizes with significant range retractions resulting in many at risk populations (Festa-Bianchet, Ray, Boutin, Côté, & Gunn, 2011; COSEWIC, 2014-2017). We demonstrate the importance of investigating historical, pre-LGM, demographic patterns to fully reconstruct the origin of the diversity within a species, which is now becoming easier with less costly sequencing and the development of programs such as PSMC. We also find that reconstructing historical events may also help us to predict how a species will respond under future climate changes, and, importantly, whether the response may vary within a species given intra-specific variation, such as between caribou ecotypes.

## ACKNOWLEDGEMENTS

Funding was provided through an NSERC Collaborative Research & Development (CRD) grant, NSERC grant RGPIN-2015-04477, Manitoba Hydro, Saskatchewan Power and Weyerhaeuser Inc. We would like to thank all the field workers for carefully collecting and storing samples from the field, Marina Kerr, Jill Lalor, Bridget Redquest, and Austin Thompson for technical support in the laboratory and Sonesinh Keobouasone for generating maps and aiding with data management. We are thankful to Ella Clark for help with literature searches. We are also thankful to the facilities of the Shared Hierarchical Academic Research Computing Network (SHARCNET: www.sharcnet.ca) and Compute Canada/ Calcul Canada gme-665-ab.Compute Canada (RRG gme-665-ab), and Amazon Cloud Computing for high-performance computing services.

## AUTHOR CONTRIBUTIONS

R.S.T. carried out analyses of genomes and wrote the manuscript. M.M. conceived the study, secured funding, and edited the manuscript. C.F.C.K. helped to conceive the study and write the manuscript. J.L.P., A.S., D.H., A.K., N.C.L., M.G., and H.S. coordinated or collected samples and edited the manuscript. P.J.W. conceived the study, secured funding, did the mitochondrial DNA analysis, and edited the manuscript.

## DATA AVAILABILITY STATEMENT

Raw reads for 18 genomes are available on the National Centre for Biotechnology (NCBI) under BioProject accession number PRJNA634908. Data will be made available on for the other genomes on the NCBI upon acceptance. Mitochondrial sequence data not already available on GenBank will also be added upon acceptance.

