## Supplementary Figures 1-8 for "Major diversification of caribou shaped by glacial cycles before the Last Glacial Maximum"

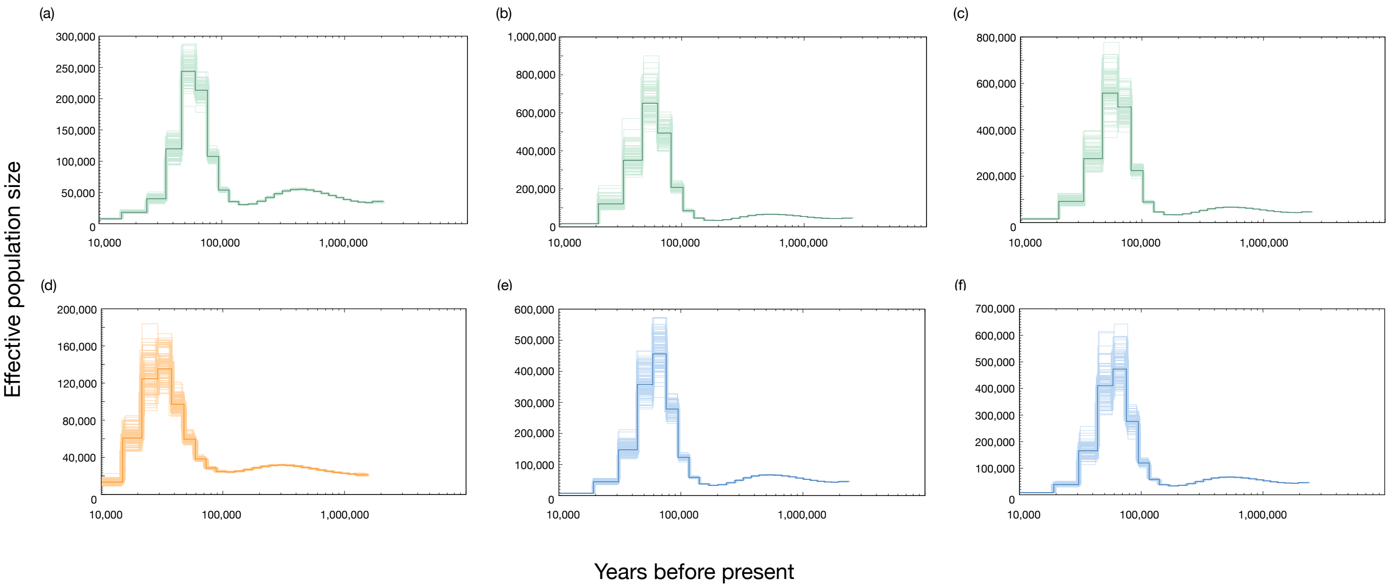


**Figure S1.** PSMC reconstruction of historical population sizes with 100 bootstrap replicates of the boreal Cold Lake (a), boreal Northwest Territories (b and c), central mountain (d), and southern mountain (e and f) caribou.

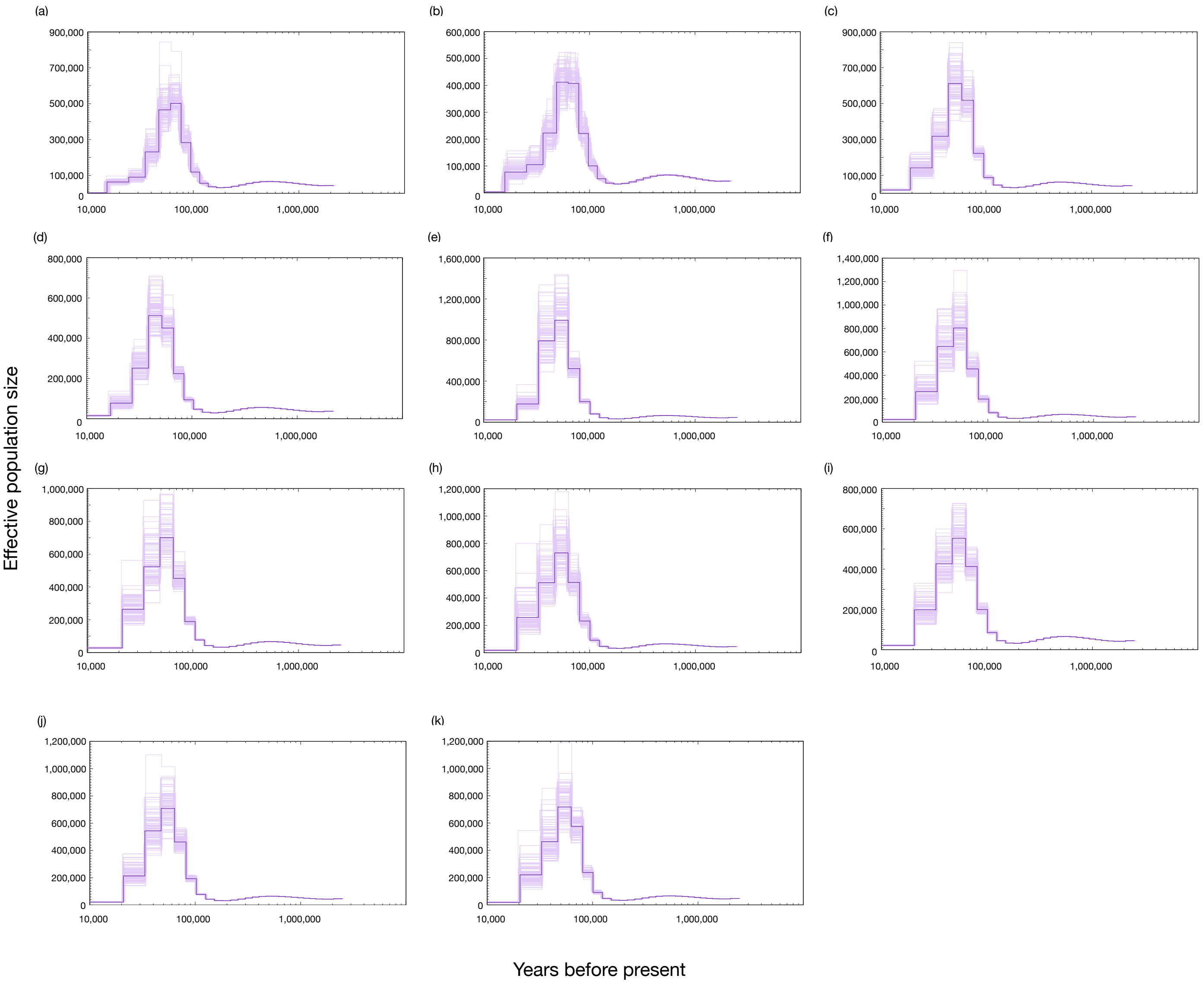


**Figure S2.** PSMC reconstruction of historical population sizes with 100 bootstrap replicates of northern mountain caribou from B.C; Itcha-Ilgachuz (a and b), Chase (c), Pink Mountain (d), Spatzizi (e), Tsenaglode (f), Frog (g and h), Muskwa (i), Atlin (j and k).

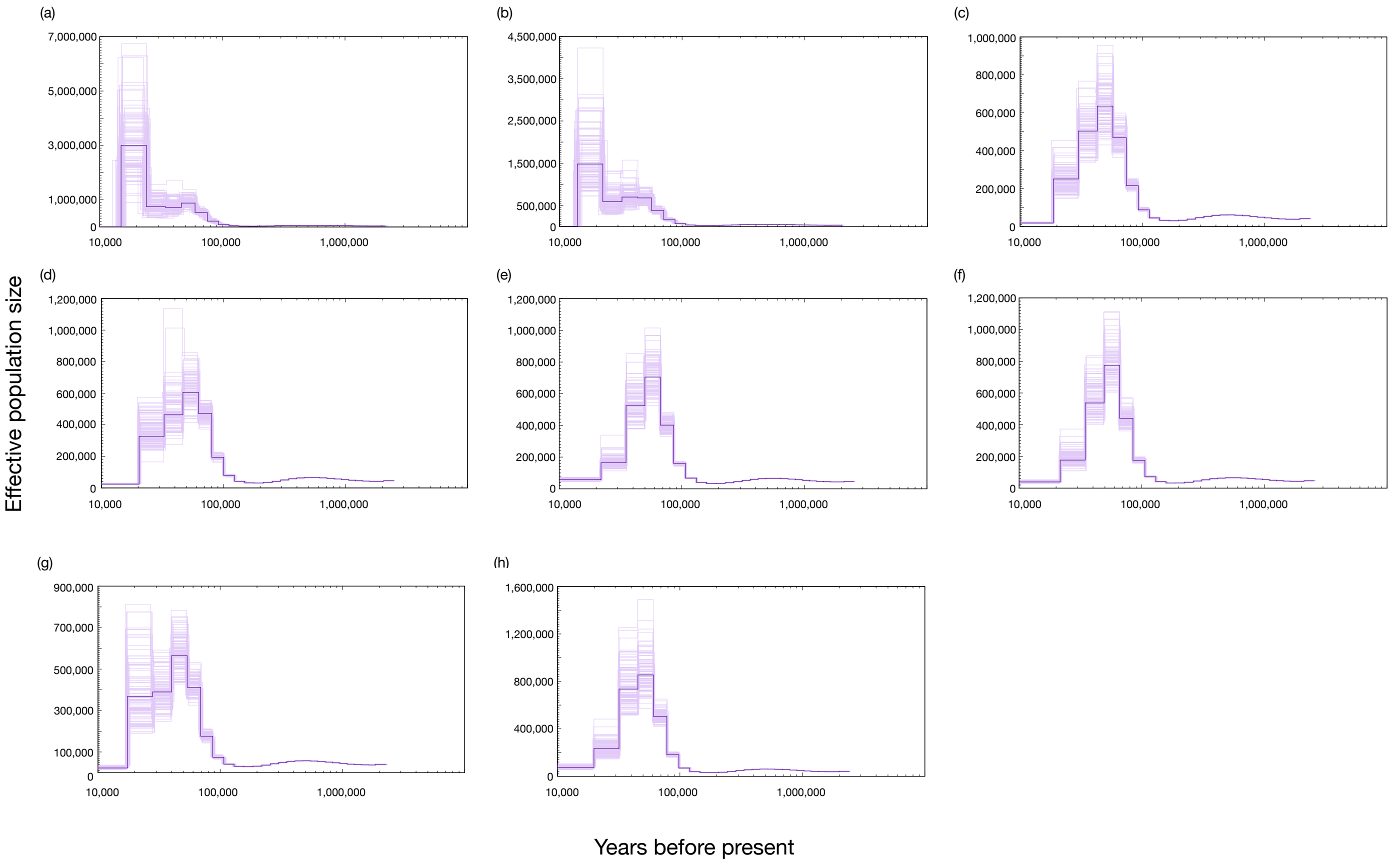


**Figure S3.** PSMC reconstruction of historical population sizes with 100 bootstrap replicates of northern mountain caribou from the Yukon and the Northwest Territories; Aishihik (a and b), Tay (c and d), Northwest Territories Redstone (e and f), Hart River (g and h).

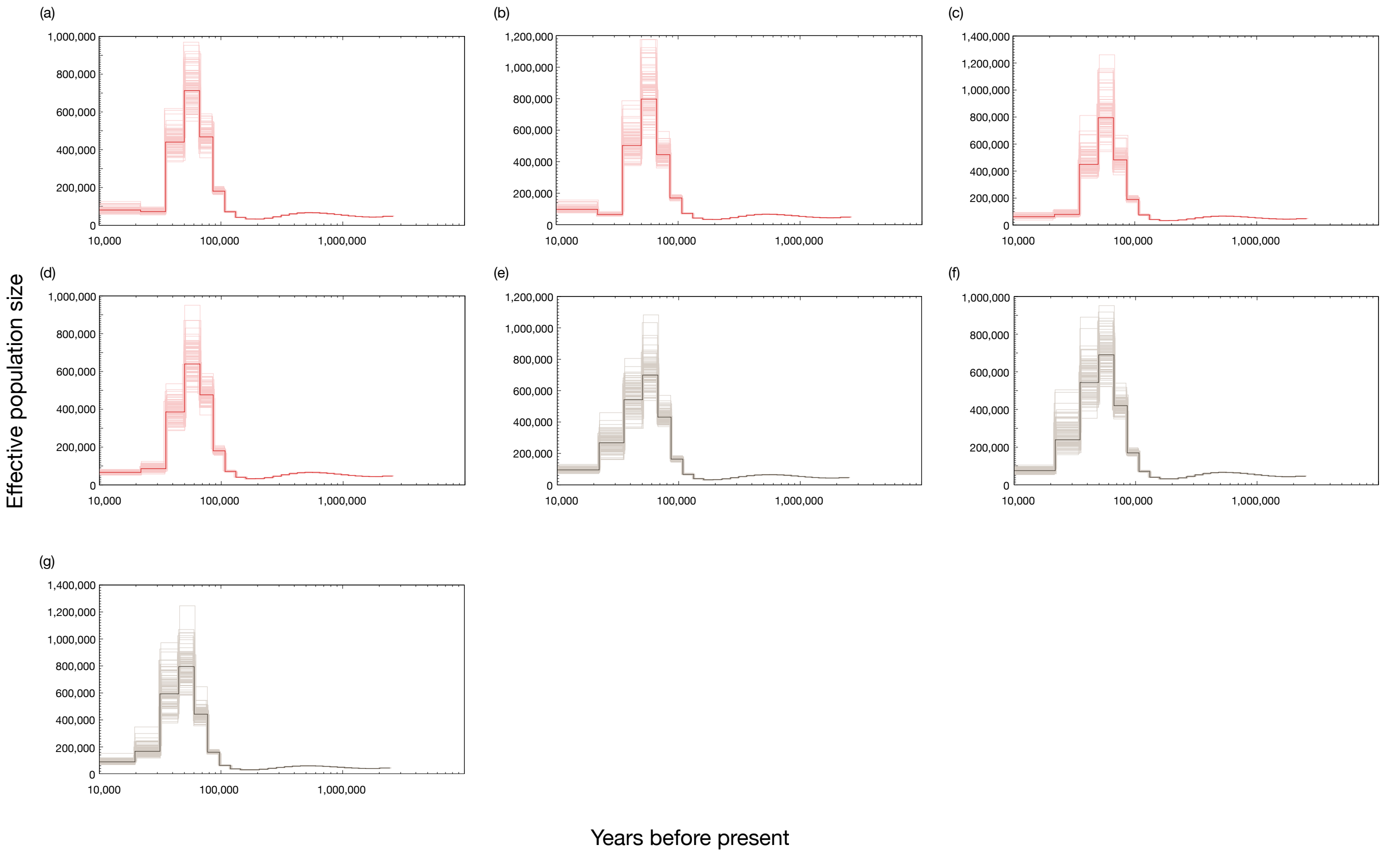


**Figure S4.** PSMC reconstruction of historical population sizes with 100 bootstrap replicates of barrenground Qamanirijuaq (a and b) and Northwest Territories Bluenose (c and d), and Grant's Porcupine (e and f) and Fortymile (g) caribou.

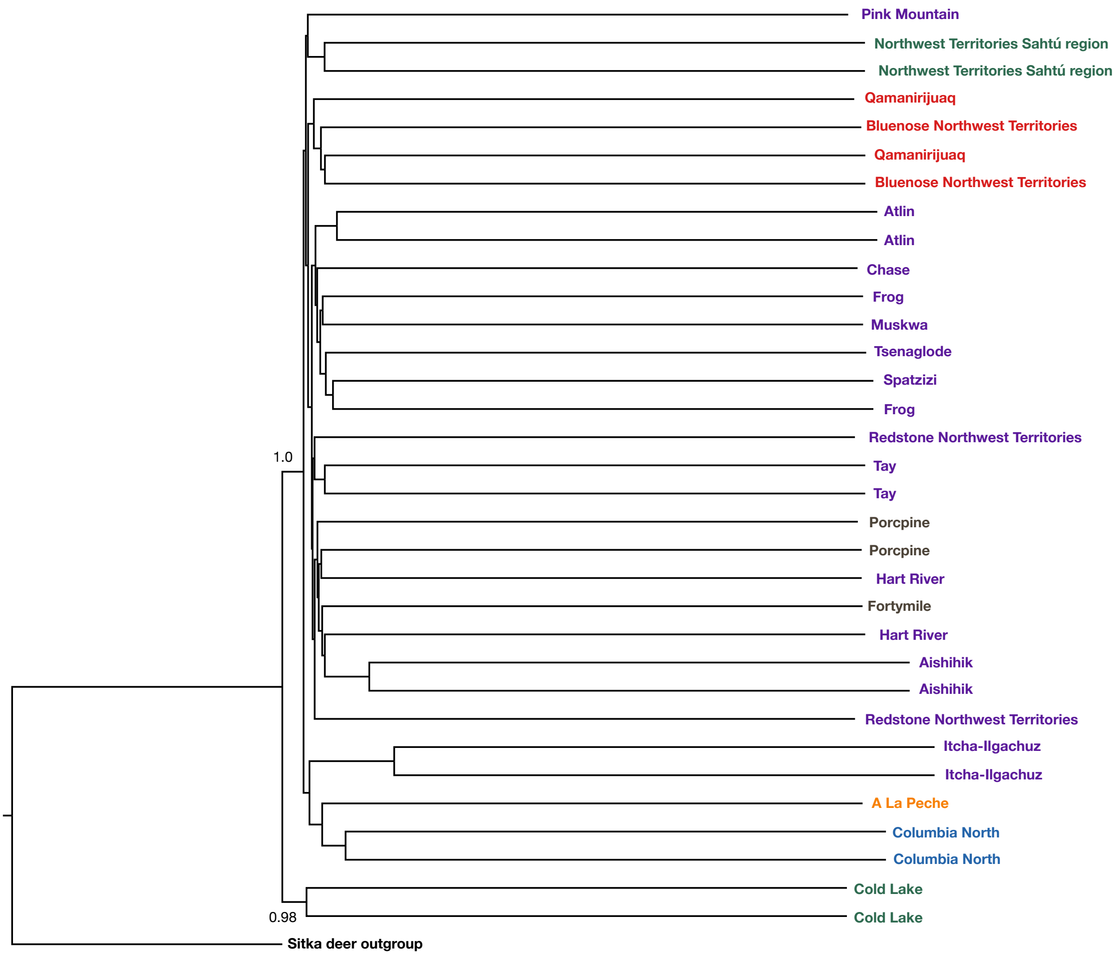


**Figure S5.** Maximum likelihood phylogenomic reconstruction from BUSCO conserved mammalian genes, with a Sitka deer outgroup. Nodes show bootstrap support values.

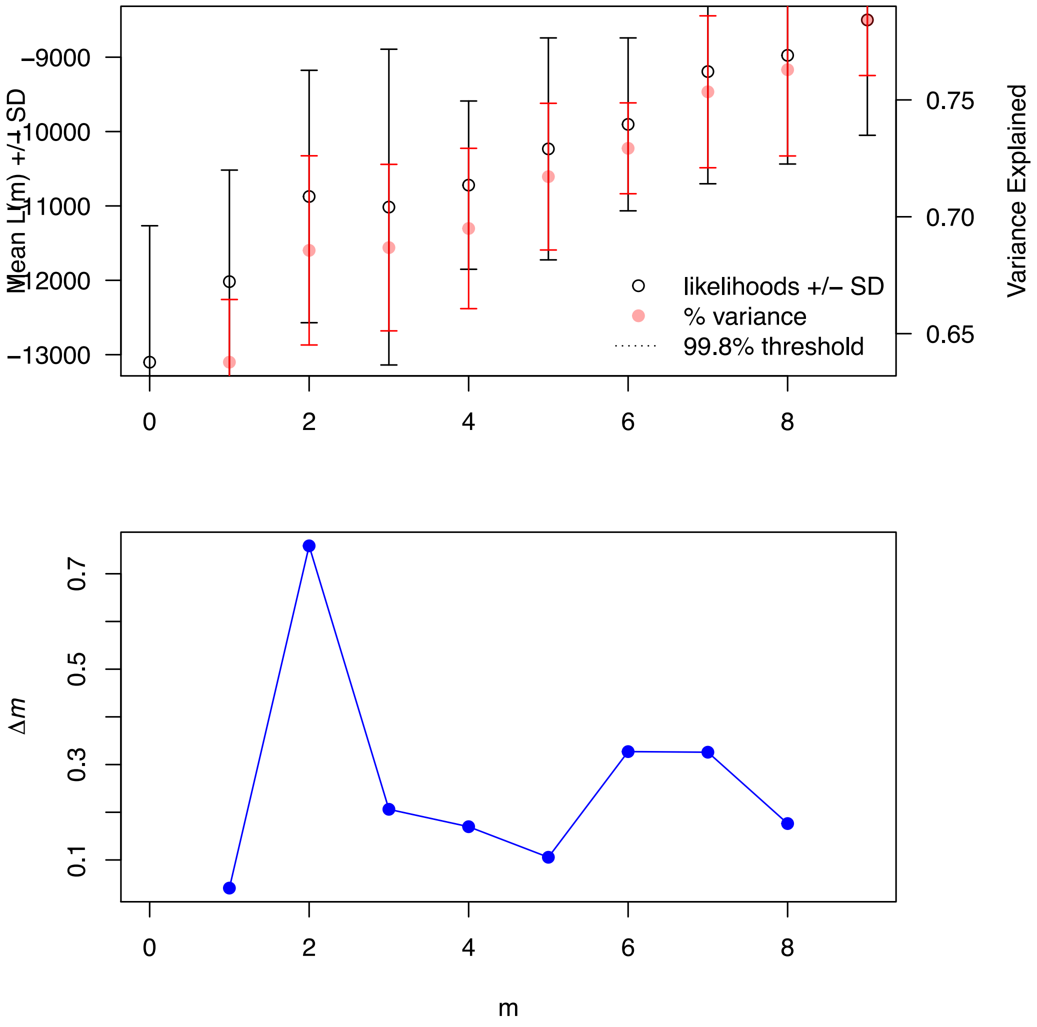


**Figure S6.** OptM results for Treemix when run for 0-9 migration events. The top plot shows the likelihood (white) and the variance explained (red) of each number of migration events. The bottom plot shows deltaM, which shows the highest peak for 2 migration events.

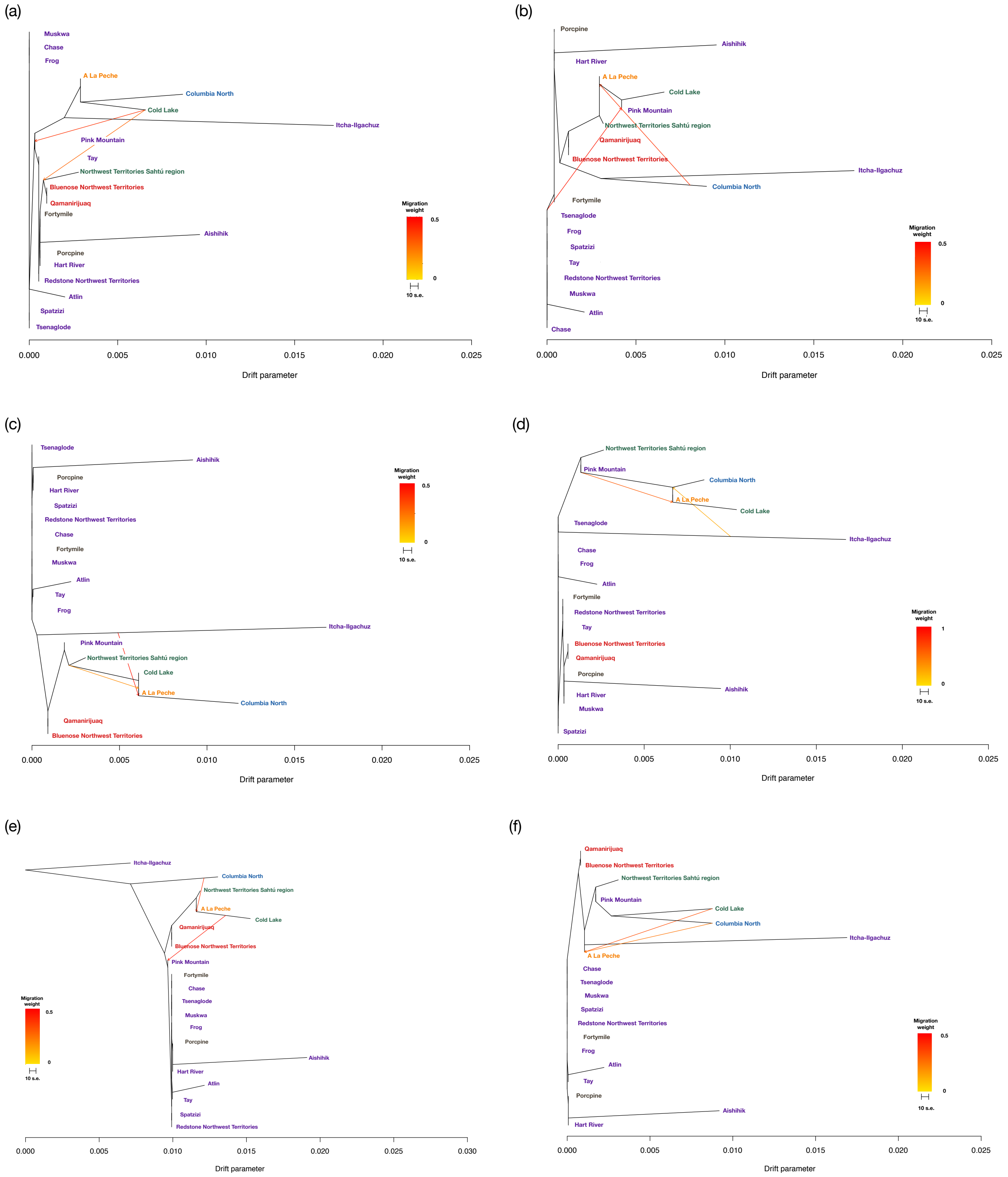

(a)
Muskwa
Chase
Frog
A La Peche
Columbia North
Cold Lake
Itcha-Ilgachuz
Pink Mountain
Tay
Northwest Territories Sahtú region
Bluenose Northwest Territories
Qamanirijuaq
Fortymile
Aishihik
Porcpine
Hart River
Redstone Northwest Territories
Atlin
Spatzizi
Tsenaglode
Migration weight
0.5
0
10 s.e.
0.000
0.005
0.010
0.015
0.020
0.025
Drift parameter
(b)
Porcpine
Aishihik
Hart River
A La Peche
Cold Lake
Pink Mountain
Northwest Territories Sahtú region
Qamanirijuaq
Bluenose Northwest Territories
Itcha-Ilgachuz
Columbia North
Fortymile
Tsenaglode
Frog
Spatzizi
Tay
Redstone Northwest Territories
Muskwa
Atlin
Chase
Migration weight
0.5
0
10 s.e.
0.000
0.005
0.010
0.015
0.020
0.025
Drift parameter
(c)
Tsenaglode
Aishihik
Porcpine
Hart River
Spatzizi
Redstone Northwest Territories
Chase
Fortymile
Muskwa
Atlin
Tay
Frog
Itcha-Ilgachuz
Pink Mountain
Northwest Territories Sahtú region
Cold Lake
A La Peche
Columbia North
Qamanirijuaq
Bluenose Northwest Territories
Migration weight
0.5
0
10 s.e.
0.000
0.005
0.010
0.015
0.020
0.025
Drift parameter
(d)
Northwest Territories Sahtú region
Pink Mountain
Columbia North
A La Peche
Cold Lake
Tsenaglode
Itcha-Ilgachuz
Chase
Frog
Atlin
Fortymile
Redstone Northwest Territories
Tay
Bluenose Northwest Territories
Qamanirijuaq
Porcpine
Aishihik
Hart River
Muskwa
Spatzizi
Migration weight
1
0
10 s.e.
0.000
0.005
0.010
0.015
0.020
0.025
Drift parameter
(e)
Itcha-Ilgachuz
Columbia North
Northwest Territories Sahtú region
A La Peche
Cold Lake
Qamanirijuaq
Bluenose Northwest Territories
Pink Mountain
Fortymile
Chase
Tsenaglode
Muskwa
Frog
Porcpine
Aishihik
Hart River
Atlin
Tay
Spatzizi
Redstone Northwest Territories
Migration weight
0.5
0
10 s.e.
0.000
0.005
0.010
0.015
0.020
0.025
0.030
Drift parameter
(f)
Qamanirijuaq
Bluenose Northwest Territories
Northwest Territories Sahtú region
Pink Mountain
Cold Lake
Columbia North
Itcha-Ilgachuz
A La Peche
Chase
Tsenaglode
Muskwa
Spatzizi
Redstone Northwest Territories
Fortymile
Frog
Atlin
Tay
Porcpine
Aishihik
Hart River
Migration weight
0.5
0
10 s.e.
0.000
0.005
0.010
0.015
0.020
0.025
Drift parameter

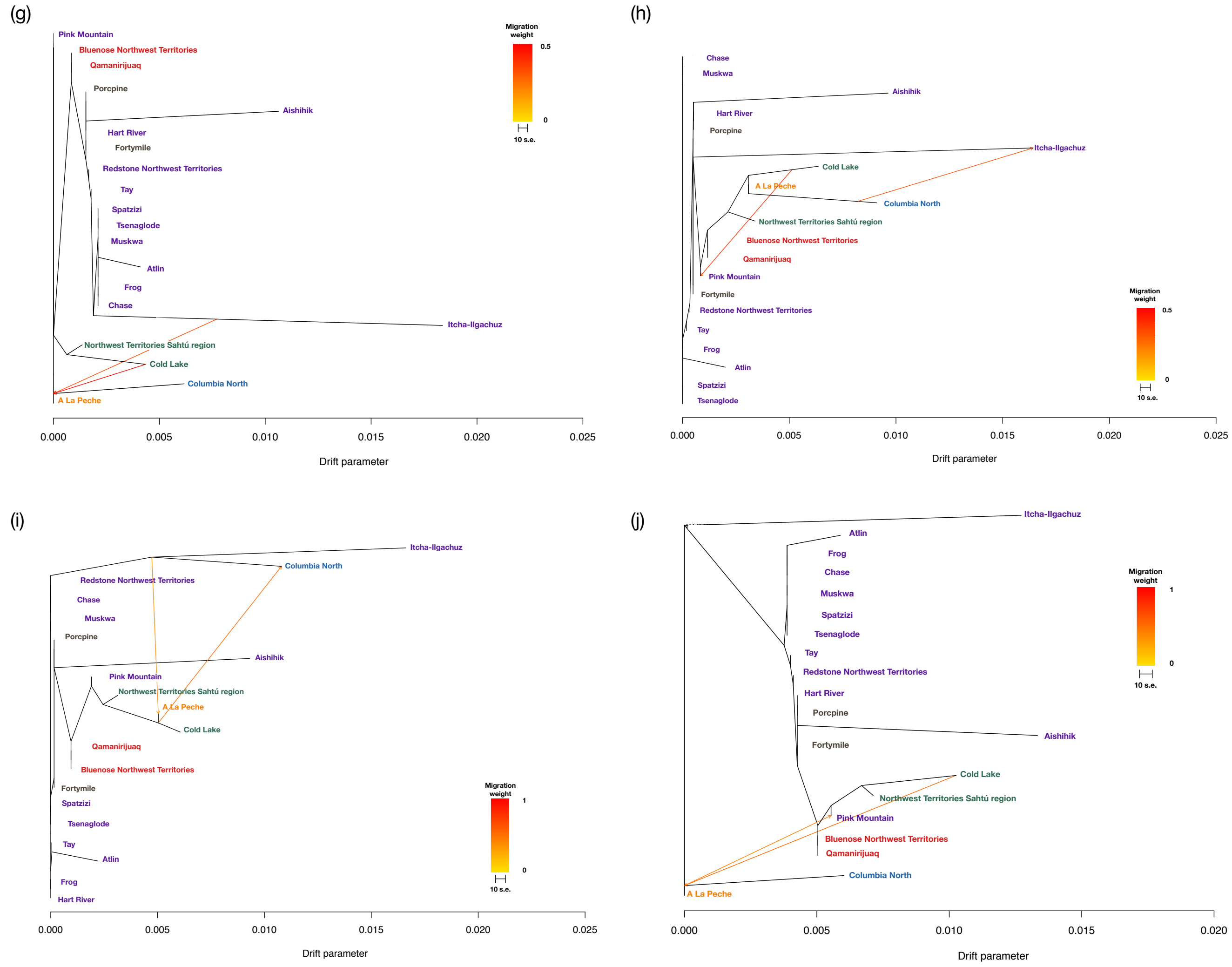


**Figure S7.** Unrooted maximum likelihood phylogenies from Treemix showing two migration events; (a) Cold Lake into both Pink Mountain and the ancestor of Northwest Territories boreal and barrenground caribou, (b) Columbia North into the ancestor of A La Peche, Cold Lake, and Pink Mountain and an ancestor of the major northern mountain clade into Pink Mountain, (c) Itcha-Ilgachuz into the ancestor of A La Peche and Columbia North, and the ancestor of Cold Lake, A La Peche, and Columbia North into A La Peche, (d) Itcha-Ilgachuz into the ancestor of Columbia North and A La Peche and Pink Mountain into A La Peche and Cold Lake, (e) Columbia North into the ancestor of A La Peche and Cold Lake and Cold Lake into Pink Mountain, (f) Cold Lake into A La Peche and Columbia North into A La Peche, (g) Itcha-Ilgachuz into the ancestor of A La Peche and Columbia North, and Cold Lake into the ancestor of A La Peche and Columbia North, (h) Cold Lake into Pink Mountain and Colmbia North into Itcha-Ilgachuz, (i) Cold Lake into Columbia North, and Columbia North into A La Peche, and (j) Cold Lake into the ancestor of A La Peche and Columbia North, and A La Peche into Pink Mountain.

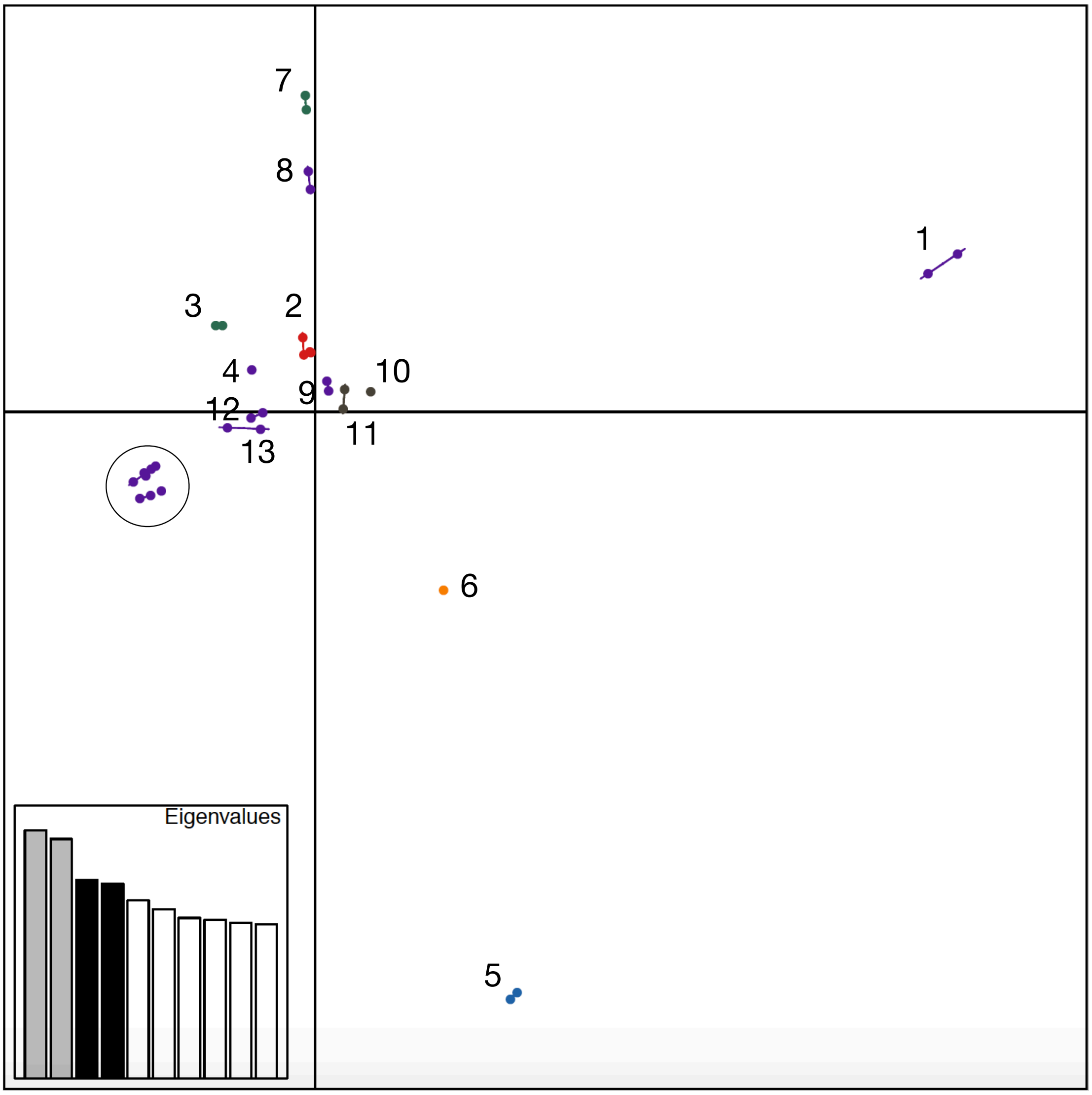


**Figure S8.** PCA plot showing the third and fourth axes, with all caribou genomes. 1. Northern mountain Aishihik; 2. All four barrenground caribou; 3. Boreal caribou from the Northwest Territories; 4. Northern mountain Pink Mountain; 5. Southern mountain Columbia North; 6. Central mountain A La Peche; 7. Boreal Cold Lake; 8. Northern mountain Itcha-Ilgachuz; 9. Northern mountain Hart River; 10. Grant's Fortymile; 11. Grant's Porcupine; 12. Northern mountain Redstone Northwest Territories; 13. Northern mountain Tay. All other northern mountain caribou are within the circle.
